# Multiple task-demands flexibly optimize neural geometry in human ventral temporal cortex

**DOI:** 10.64898/2025.11.30.691163

**Authors:** Tarana Nigam, Andrea F. Campos-Pérez, Pierre Mégevand, Juan R. Vidal, Marcela Perrone-Bertolotti, Philippe Kahane, Thomas Thesen, Orrin Devinsky, Lucia Melloni, Caspar M. Schwiedrzik

## Abstract

A hallmark of human intelligence is the ability to perform multiple tasks immediately upon instruction. Yet, the neural processes that implement such flexibility remain unclear. Using intracranial electrophysiology in epilepsy patients, we examined how representational geometries evolve as participants switched among three tasks - individuation, categorization, and conceptualization - on a trial-by-trial basis. Although tasks differed in the required representational geometries, neural representations across cortex were not immediately optimized following task cues. Instead, task-tailored geometries emerged gradually over successive stimulus repetitions within a trial. Ventral temporal cortex was the only region to exhibit task-specific adjustments for all three tasks, dynamically transforming within- and between-category distances in line with task demands. Importantly, these gradual representational changes were behaviorally relevant, predicting trial-level performance. Our results show that behavioral flexibility is supported by incremental, task-dependent refinement of representational geometry already within sensory cortex - far earlier in the processing hierarchy than previously thought.

## Introduction

A hallmark of human intelligence is our ability to perform multiple, including entirely new tasks - even when based solely on verbal instructions^1,2^. Such flexibility is critical for adapting to novel situations and changes in the environment, while deficits in this domain are observed in many neuropsychiatric disorders. Rapid flexibility is thought to rely on the reuse of information from previously learnt tasks, i.e., compositionality^3–5^. Yet, how this potentially uniquely human capacity for flexibility comes about on a neural level remains a central unanswered question.

A critical determinant of such flexible reuse is the geometry of the underlying neural representations, i.e., the degree of overlap and separation of neural activity patterns^6,7^: small distances between activity patterns in a high-dimensional activity space enable grouping and generalization, while larger distances reflect distinctions and hence discriminability. To achieve an optimal representational geometry for the task at hand, while minimizing interference with other tasks, the brain could reconfigure representations based on task demands^8,9^. Such reconfiguration may manifest as a task-dependent expansion of neural distances along behaviorally relevant feature dimensions and a compression along irrelevant ones, thereby producing task-tailored representational geometries. Thus, by adjusting representational geometry, multiple different tasks can be performed on the same set of inputs (Fig. 1A).

**Figure 1:**
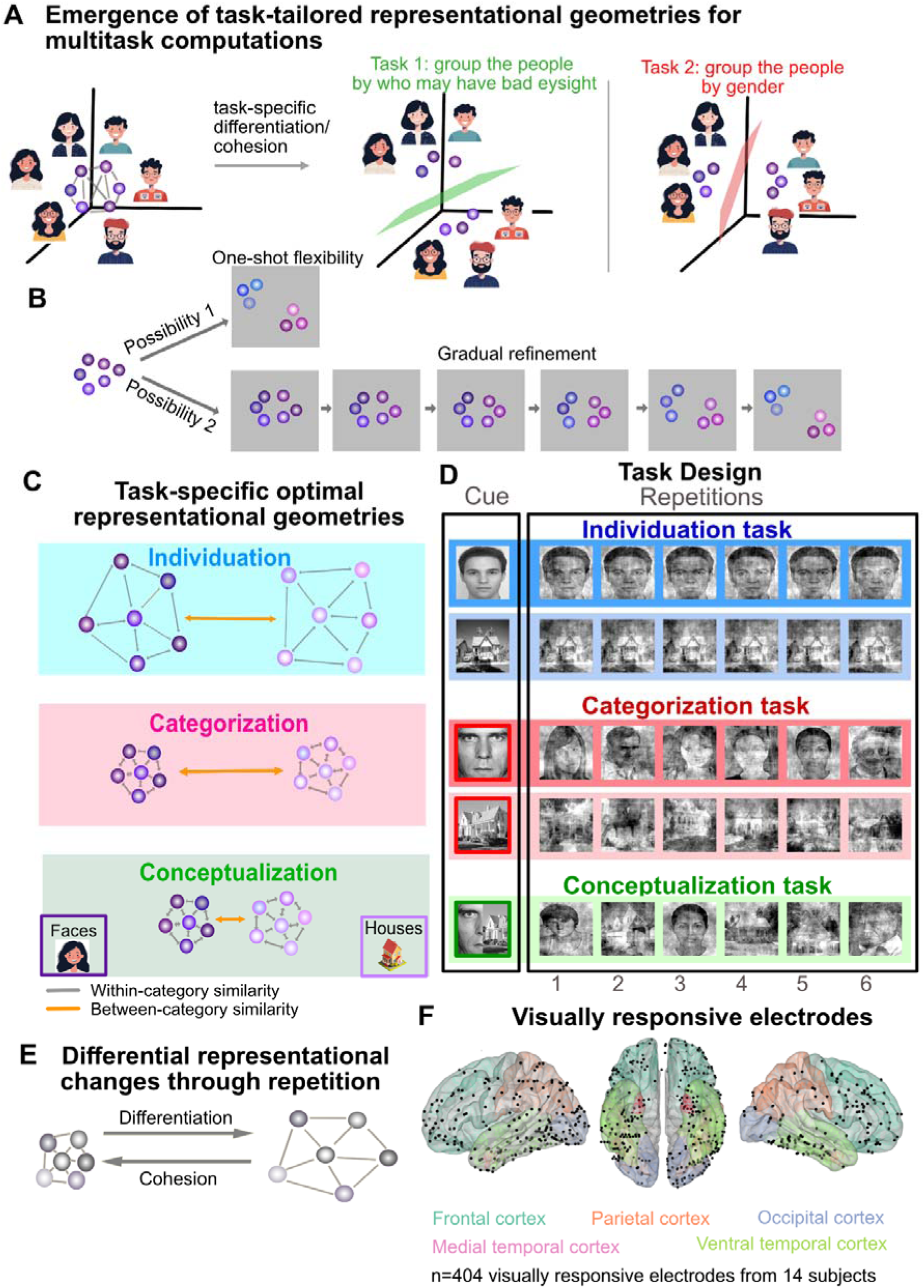
Hypotheses and Task Design. **A:** Flexible behavior depends on the format of how information is represented in neural population activity. The relationships among neural activity patterns for different stimuli or tasks define the representational geometry. Humans can rapidly perform novel tasks, e.g., grouping faces based on who may have bad eyesight. One way to solve this is by expanding task-relevant and compressing irrelevant dimensions, transforming representations into task-specific geometries. Flexibility in compressing or expanding allows performing multiple tasks on the same stimuli, e.g., grouping the same faces based on gender. **B:** Competing hypotheses for how rapidly flexible task representations emerge in the human brain. *Possibility 1:* Humans exhibit zero- or one-shot flexibility, where representations reconfigure into task-optimal geometries immediately upon instruction*. Possibility 2:* Representations evolve gradually through repeated experience. **C:** Hypothesized task-tailored representations across the three tasks in this study: the individuation task requires differentiation of neural representations both within (grey arrows) and between (yellow arrows) visual categories (faces, houses). The categorization task requires cohesion within each category and differentiation between categories. The ad hoc conceptualization task requires cohesion within each category and across two perceptually distinct categories. **D:** Subjects flexibly performed three tasks: individuation, categorization, and conceptualization. These tasks were presented in the framework of an oddball paradigm. The subjects were instructed to report after every trial whether a deviant (28.75% of trials) had occurred. Oddballs could occur on the repetition 1 or later with equal probability. Subjects were cued which task to perform on a trial-by-trial basis. In the individuation task, the cue was followed by a sequence of 6 images depicting the same identity (face or house). In the categorization task, the repeating images varied over repetitions but were all from the same category (either faces or houses). In the conceptualization task, the repeating images were either faces or houses, forming an ad hoc concept. Accurate deviant detection depended on the task at hand: in the individuation task, the deviant could be that of a different identity or category; in the categorization task, the deviant was outside of the cued visual category; in the conceptualization task, detecting a deviant entailed identifying an image that was neither a face nor a house (i.e., an image from a different object category). **E:** Hypothesized optimization of neural representations across repetitions. With repeated stimulus presentation, representations are expected to undergo task-dependent refinement - differentiation (increased neural distances, hence decreased pattern similarity) or cohesion (decreased neural distances, hence increased pattern similarity) in a manner optimal for performing each task. **F:** A total of 404 visually responsive electrodes from 14 epilepsy patients, identified via an independent localizer, were included in the analysis. These electrodes were distributed across five brain regions: 45 in the occipital cortex, 140 in the ventral temporal cortex, 61 in the parietal cortex, 33 in the medial temporal lobe, and 125 in the frontal cortex. Note: the axial view is displayed from the ventral side. Cartoon faces and houses in A and C are reproduced with permission from Adobe.

Behavioral evidence suggests that humans are remarkably efficient in reusing preexisting information, performing new tasks essentially one- or even zero-shot. E.g., children can infer the meaning of a completely novel word from context after a single exposure (“fast mapping”^10^), and adults can generalize abstract rules to entirely new combinations without prior examples^11^. This would suggest that the underlying neural representations should transform into task-tailored, optimal geometries instantaneously upon instruction *(Possibility 1)*. However, theoretical work has shown that distributed neural representations typical of the neocortex require repeated experience to render them non-overlapping^12,13^. Gradual changes in representational geometries are moreover beneficial in reducing and smoothing noise such that a few, idiosyncratic and/or atypical experiences do not disrupt existing representations. This implies that representations may transform into task-specific geometries only more slowly, through gradual experience *(Possibility 2)* (Fig. 1B).

Here, we ask how flexibly and how rapidly the human brain implements representational transformations for performing multiple tasks. To this end, we conducted invasive electrophysiology across a wide range of brain areas in humans undergoing epilepsy monitoring while they performed three different tasks using the same set of complex visual stimuli. These tasks (i) required distinct representational geometries to be solved, (ii) were cued on a trial-by-trial basis, demanding frequent switches between geometries, and (iii) were designed to reveal whether the appropriate geometry is implemented immediately upon instruction or gradually refined with experience throughout a trial.

The three tasks entailed i) *individuation* (e.g., distinguishing a face from other faces), ii) *categorization* (e.g., distinguishing faces from houses), and iii) *ad hoc conceptualization* beyond perceptual similarities (grouping faces and houses into one ad hoc concept/category, to be distinguished from other objects), respectively. The tasks were carried out in the framework of an oddball paradigm^14^ (Fig. 1D): on each trial, the task cue was followed by a sequence of six images and subjects had to report whether or not a deviant stimulus occurred. Whether a stimulus qualified as a deviant depended on the task at hand: in the *individuation* task, the repeating stimulus was a specific face or house identity and any change in identity counted as a deviant; in the *categorization* task, the repeating stimuli belonged to a visual category (faces or houses) and any stimulus from a different category counted as a deviant; and in the *ad hoc conceptualization* task, the repeating stimuli were faces or houses and any non-face, non-house object qualified as a deviant.

To successfully identify a deviant in each of the three tasks, the existing representational geometry should be differentially transformed into task-tailored geometries through differentiation and/or cohesion: e.g., to distinguish a deviant from standard faces in the individuation task, within category distances should be higher than in the categorization and conceptualization tasks; conversely, to form an ad hoc concept/category^15^ grouping faces and houses and to distinguish it from other objects in the conceptualization task, between category distances between faces and houses should be smaller than in the individuation and categorization tasks (Fig. 1C). Because the deviant could already occur as the first stimulus within a trial (repetition 1), any task-specific optimization could be present immediately following the cue, i.e., zero- or one-shot. However, if task-tailored differentiation or cohesion emerged only over several repetitions within a trial, our design allowed us to track such gradual, within-trial changes in representational geometry (Fig. 1E).

We examined the flexibility of representational geometries in frontal and parietal cortex, sensory areas (occipital and ventral temporal cortex), and the medial temporal lobe. Surprisingly, none of these regions exhibited task-tailored representations upon cueing. Instead, we find that representations gradually evolved within a trial towards task-tailored configurations. Interestingly, the only region that exhibited task-specific optimization for all three tasks was the ventral temporal cortex, where neural representations exhibited flexible differentiation or cohesion based on task demands. Our findings thus suggest that task-tailored representational geometries underlying flexible multi-tasking develop through gradual, context-dependent optimization of multipurpose representations in sensory cortices and hence originate earlier in the cortical hierarchy than previously thought.

## Results

To examine how task-tailored representational geometries emerge in the human brain, we recorded intracranial electrophysiology from a large sample of epilepsy patients (n=14), covering a wide range of brain areas with over 2000 electrodes. Subjects were initially verbally instructed and then cued on a trial-by-trial basis to perform one of three tasks (individuation, categorization, or conceptualization) over the same set of visual stimuli (faces, houses, and other objects). These tasks were selected to probe multitasking independent of stimulus features and to allow flexible grouping based on both perceptual similarity and more abstract conceptual relations. Further, the three tasks were conducted in an oddball format, ensuring identical responses (output) for detecting deviants. At the same time, the design did not rely on preexisting or frequently repeating stimulus-response mappings, rendering every trial unique. Because the oddball stimulus could appear as the first of six repetitions or later, we could assess if and how quickly neural representations adapted to different tasks.

All subjects performed with above-chance accuracy (mean=81.20%, standard error of mean=±2.02%, range=65.71-94.29%). Performance on the different tasks did not differ statistically in terms of accuracy (repeated measures ANOVA, F(4,52)=2.0973, p=0.0944) or reaction times (RT; repeated measures ANOVA, F(4,52)=1.7232, p=0.1589), indicating that the difficulty of the three tasks was well matched. Accuracy to detect a deviant was above chance (but not perfect) already for deviants occurring on the first repetition (overall mean accuracy=86%) and did not differ significantly between tasks (ANOVA, accuracy: F(2,22)=0.4493, p=0.6438; RT: F(2,20)=F=1.7827, p=0.1939; mean accuracy individuation task=88.13%, mean accuracy individuation task=86.641%, mean accuracy conceptualization task=93.33%).

To assess if and how neural representational geometry is differently modified across tasks, we initially focus on trials in which the subjects correctly performed the task and images were repeated without an intervening deviant (71.25% of all trials). Since all three tasks involve visual stimuli, in our main analyses, we investigate 404 visually responsive electrodes (Supplementary Fig. 1 – Upper Panel) as determined by an independent localizer. These were found in occipital cortex (45 electrodes), ventral temporal cortex (140 electrodes), parietal cortex (61 electrodes), the medial temporal lobe (33 electrodes), and frontal cortex (125 electrodes) (Fig. 1F), and at least 8 of the 14 subjects contributed to each of the clusters (Supplementary Table 1 and Supplementary Fig. 1 – Lower Panel). We consider the high-gamma frequency band (HGB, 70-200Hz) which correlates with multiunit firing rate^16,17^.

To assess representational geometries, we determine the similarity between stimulus-evoked neural time courses (both across the individual stimuli within a category as well as between categories) for each electrode^18,19^ (Fig. 2A). This approach captures a fundamental aspect of representational structure - the temporal evolution of neural responses, and reflects the computation of temporal pattern separation^18–20^. Previous work has shown that temporal response profiles provide a rich source of stimulus-specific information in both cortex^21,22^ and hippocampus^19^, where temporal codes carry discriminative information in addition to response magnitude^23,24^. By comparing the full temporal dynamics of neural activity, we thus account for representational distinctions that unfold over time. We also empirically validated that this measure captures a meaningful dimension of stimulus representation (see Supplementary Section 1). In addition, we also estimated representational structure from the similarity of spatial activity patterns across electrodes within each ROI, capturing population-level information distributed across electrodes (see Supplementary Section 2).

**Fig. 2:**
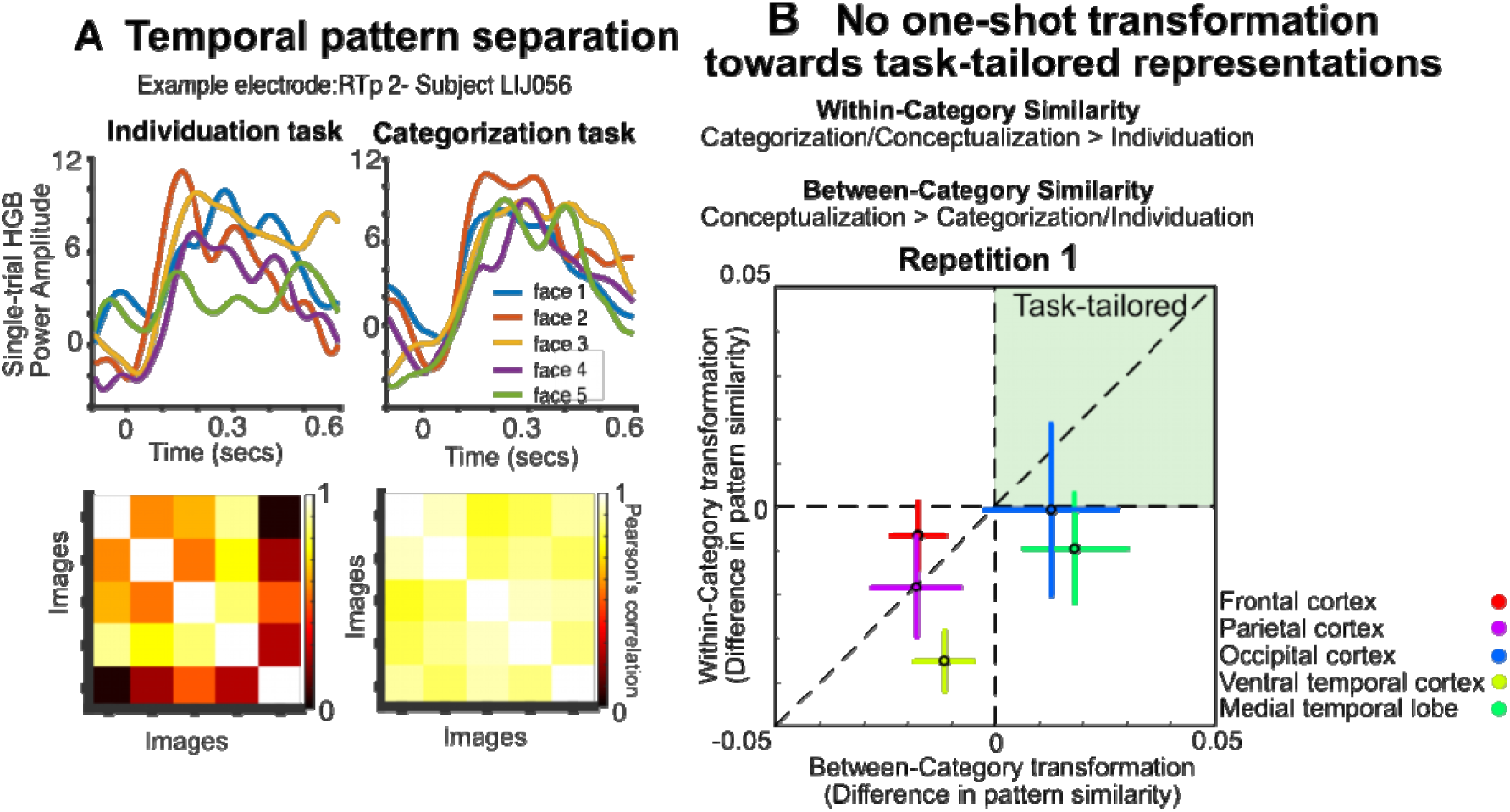
Analysis approach and result for task-tailored representation on repetition 1. **A.:** We assessed neural representations by analyzing the temporal dynamics of single-trial responses. For each task and repetition, we computed the similarity between neural time courses in response to all presented images. As an example, we show an electrode from ventral temporal cortex (RTp2, subject LIJ056). The upper panel displays single-trial high-gamma responses for five images in repetition 1 of the individuation (left) and categorization (right) tasks. The lower panel shows the representational similarity matrix derived from pairwise time-course similarities, with warm colors indicating higher similarity. Similarity values were then averaged across images within and between categories. **B:** Task-tailored representational transformations at the first repetition. The 2-D plot shows differences in pattern similarity: the x-axis represents within-category similarity and the y-axis represents between-category similarity. Task-tailored areas are expected in the upper-right quadrant, showing increased within-category and between-category similarity consistent with task demands; all other quadrants reflect suboptimal representations that are not task-tailored. Specific contrasts to assess task-tailored representations: within-category similarities in categorization and conceptualization task > individuation task and between-category similarities in conceptualization task higher in conceptualization task> categorization and individuation task. No brain area showed statistically significant task-tailored representational change in the temporal pattern of responses (one-sided t-tests on within and between category pattern similarity changes against 0, all p > 0.075). See Supplementary Fig. S5 for results from the spatial pattern of responses and Supplementary Section 5 for an analysis including all, not only visually responsive electrodes.

### Representational transformations do not occur instantaneously

To determine whether the brain implements task-tailored representational geometries as soon as a new task is cued, we examined representational similarities at the first stimulus repetition within each trial. Each of the three tasks has a specific geometry of within and between category distances that could support optimal task performance: the individuation task demands differentiation of neural representations within and between categories; the categorization task requires cohesion within each category; and the conceptualization task requires both within and between-category cohesion. Hence, for a representation to be task-tailored, there should be higher within-category similarity in the categorization and conceptualization tasks as compared to the individuation task, and between-category similarity should be higher in the conceptualization task than in the individuation and categorization tasks. To assess whether within and between category similarities jointly transformed towards task-tailored representations according to these predictions, we computed differences in both types of similarity between the tasks. Surprisingly, we find that representations in none of five areas under investigation are demonstrably task-tailored at repetition 1, i.e., none of the areas show statistically significant differences in temporal pattern similarity greater than 0 (Fig. 2B; change in within and between category transformation, one-sided t-test against 0: all p-values>0.075).

As a complementary analysis, we also computed specific contrasts that define task-tailored representations for the three tasks, and used a combination test to determine whether within and between category similarities jointly transformed towards the optimal geometries (within category similarity: Categorization>Individuation and Conceptualization>Individuation; between category similarity: Conceptualization>Categorization and Conceptualization>Individuation). We again find that on repetition 1, representations in none of the five brain areas show any combined difference between the tasks (Stouffer’s combination test; Frontal cortex: test statistic: -2.865, combined p=1.000; Parietal cortex: test statistic: -2.963, combined p=1.000; Occipital cortex: test statistic: 0.821, combined p=0.9345; Ventral temporal cortex: test statistic: -5.255, combined p=1.000; Medial temporal lobe: test statistic: 0.89, combined p=0.9345; p-values corrected for multiple comparisons).

Together, this suggests that even though subjects can perform all three tasks above chance even when the oddball stimulus occurs at the first repetition, the underlying representations do not exhibit task-tailored transformations immediately upon instruction.

### Task-tailored geometries emerge gradually

Given that task-tailored representations did not appear already at the outset of a trial, we further investigated whether they gradually emerged on task, as would be suggested by theory^12,25^. Indeed, behaviorally, subjects correctly detected deviants faster if they were exposed to a greater number of repetitions in a trial (in a quadratic fashion, F(1,60)=9.653, p=0.00289). This indicates that repetition indeed leads to continual optimization for solving the tasks and that prior information is being used to become faster with repeated exposure.

To determine whether transformations towards task-tailored optimal neural geometries occur over repetition within a trial, and if so, where in the brain that happens, we assessed whether similarities in temporal response profiles varied with task and repetition differentially between brain areas. Indeed, when combining all data, we find that different tasks undergo distinct patterns of neural optimization across the different brain areas for within and between category similarities (region × task × repetition interaction; within category similarities: F(40, 12025)=1.6916, p=0.0041, partial eta squared = 0.0056; between category similarities: F(40, 6188)=1.6413, p=0.0066, partial eta squared = 0.0105). To determine the individual brain areas’ contributions to on task optimization, we proceed to investigate the representational transformation over the repetition in the frontal-parietal network, sensory areas, i.e., occipital and ventral temporal cortex, and the medial temporal lobe. As in the preceding analyses, we hypothesize that for representational geometries to become task-tailored through repetition, they should gradually differentiate in the individuation task, show increased cohesion within category in the categorization task, and both within and between category in the conceptualization task (Fig. 3A).

**Figure 3:**
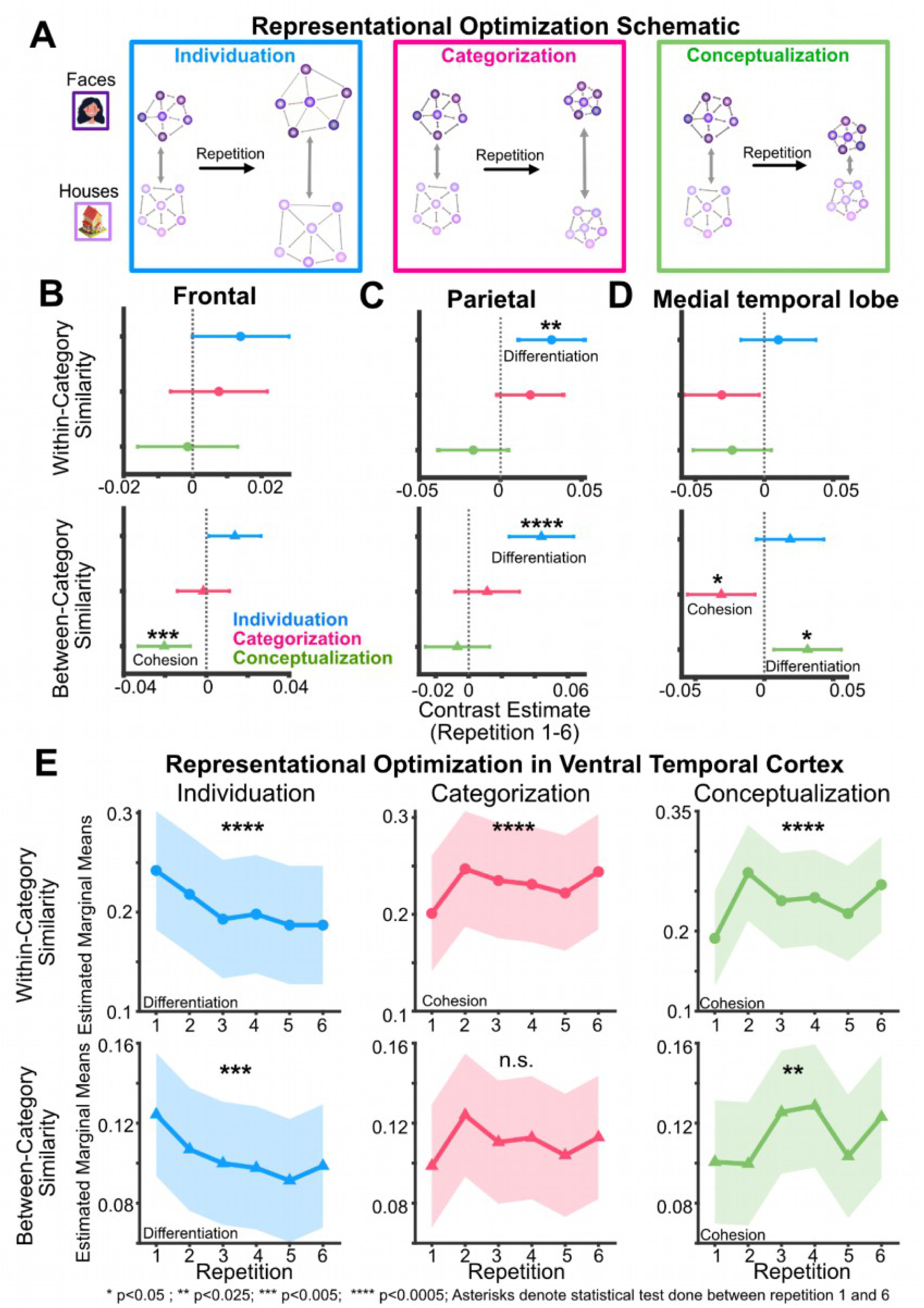
Flexible representational optimization in multiple tasks. **A.** Representational optimization: we hypothesize that repetition leads to task-specific optimization of neural representations. In the individuation task, within-category and between-category similarities decrease (differentiation of the individual stimuli both within and between visual categories (faces and houses)); in the categorization task, within-category similarity increase while between category similarity decreases, i.e., individual stimuli within a category become more similar through cohesion of neural representations within a category and different categories (faces and houses) more distinct from each other; in the conceptualization task, both within-category and between-category similarity increases with repetition, i.e., cohesion of both individual stimuli within a category as well as between stimuli of perceptually different visual categories (faces and houses). **B.** Frontal cortex: between category similarities of temporal patterns decreased with repetition in the conceptualization task (repetition 1 vs. 6, t(2023)=-3.158, p=0.0048); while no statistically significant representational optimization occurred for any of the other tasks (all p>0.06). All p-values are corrected for multiple comparisons for the number of tasks. The x-axis indicates contrast estimates of the post-hoc test between repetition 1 vs. 6. Error bars are model-based standard error. Positive x-values indicate differentiation over repetitions, i.e., decreasing similarity over repetitions, and negative values indicate cohesion over repetitions (applies to panels C and D also) **C.** Parietal cortex: Increasing differentiation over repetition (within and between category) was observed in the individuation task, i.e., decreasing similarity (between-category similarities: repetition 1 vs. 6, t(969)=4.411, p<0.0001; within-category similarities: repetition 1 vs. 6, t(1874)=3.04, p=0.0071). In the categorization and conceptualization tasks, we found no statistically significant evidence for optimization between categories (repetition 1 vs. 6, all p>0.52) or within categories (repetition 1 vs. 6, all p>0.14). **D.** Medial temporal lobe: No statistically significant optimization over repetitions occurred for between category similarities in the individuation task (repetition 1 vs. 6, t(510)=1.500, p=0.1341), while there was increasing between-category cohesion in the categorization task (repetition 1 vs. 6, t(510)=-2.483, p=0.0378). Increasing differentiation between categories was observed in the conceptualization task, i.e., between-category similarities decreased over repetitions (repetition 1 vs. 6, t(510)=2.504, p=0.0378) when the task demands required integration or cohesion of the visual categories over repetition. The available data showed no statistically significant task optimization in terms of within-category similarities (all p>0.07). **E.** Ventral temporal cortex: Neural representation in the ventral temporal cortex exhibit differential optimization for multiple tasks (significant interaction repetition × task for both between-category (F(10, 2244)=4.8322, p<0.0001) and within-category similarities (F(10, 3546)=7.1779, p<0.0001)). Increasing differentiation over repetition (within and between category) was observed in the individuation task, i.e., decreasing similarity (between-category similarities: repetition 1 vs. 6, t(2244)=3.167, p=0.0047; within-category similarities: repetition 1 vs. 6, t(3546)=4.65, p<0.0001). Increasing cohesion within a category over repetition was observed in the categorization task, i.e., increasing similarity (repetition 1 vs. 6, t(3546)=-3.65, p=0.0002), but did not reach statistical significance between categories (repetition 1 vs. 6, t(2244)=-1.752, p=0.0799); further cohesion increased within a category (repetition 1 vs. 6, t(3546)=-5.11, p<0.0001) and between perceptually different visual categories (repetition 1 vs. 6, t(2244)=-2.770, p=0.0113) for the ad hoc conceptualization task. The y-axis represents estimated marginal means of the within and between category similarity; error bars represent the model-based standard error. Asterisks denote statistical tests done between repetition 1 and 6 per condition; *p<0.05; **p<0.025; ***p<0.005; ****p<0.0005; p-values are corrected for multiple comparisons for the number of tasks. See Supplementary Section 2 for an analysis of the spatial pattern of responses and Supplementary Section 5 for an analysis including all, not only visually responsive electrodes. The cartoon face and house in A are reproduced with permission from Adobe.

Given the role of frontal and parietal cortex in cognitive flexibility and multitask computations^2,26,27^, we first ask whether these two areas show both flexible, task-dependent neural optimization. Starting with frontal cortex, we find that between category similarities show an interaction between task and repetition (F(10, 2023)=3.5966, p<0.0001, partial eta squared = 0.0175), indicative of task-dependent optimization. However, further inspection reveals that gradual optimization occurs only in the conceptualization task: in this task, between-category similarities decrease with repetition (Fig. 3B; repetition 1 vs. 6, t(2023)=-3.158, p=0.0048, multiple-comparisons corrected for 3 tasks, partial eta squared = 0.004906; linear trend, t(2023)=4.387, p=<0.0001, multiple-comparisons corrected for 4 trends), suggesting increasing representational cohesion between faces and houses. We found no statistically significant evidence that on-task optimization of the between-category similarities occurs for the other two tasks (Individuation: t(2023) = 2.131, p = 0.0664, partial eta squared = 0.002240, Categorization: t(2023) = -0.226, p = 0.8211, partial eta squared = 0.000025; p-values are multiple-comparisons corrected for 3 tasks). Within category similarities do not change significantly with repetition (F(10,3682.1)=1.2721, p=0.240184, partial eta squared = 0.0034). Therefore, the available evidence suggests that frontal cortex is specifically involved in joining faces and houses into one abstract concept by reducing their representational distance.

In parietal cortex, we also find an interaction of task and repetition for between category similarities (F(10,969)=2.1062, p=0.0216, partial eta squared = 0.0213), albeit with a different profile. Specifically, between category similarities change only in the individuation task, decreasing with repetition (Fig. 3C; repetition 1 vs. 6, t(969)=4.411, p<0.0001, partial eta squared = 0.019684, all other p>0.52, multiple-comparisons corrected for 3 tasks; Categorization: partial eta squared = 0.001309, ad hoc conceptualization: partial eta squared = 0.000480); linear trend, t(969)=-4.030, p=0.0002 and quadratic trend, t(969)=3.033, p=0.0075, multiple-comparisons corrected for 4 trends). Within category similarities follow the same pattern: similarities decrease with repetitions only in the individuation task (repetition 1 vs. 6, t(1874)=3.04, p=0.0071, partial eta squared = 0.004907, all other p>0.14 multiple-comparisons corrected for 3 tasks; Categorization: partial eta squared = 0.001707, ad hoc Conceptualization: partial eta squared = 0.001231). Hence, parietal cortex seems to be specifically involved in optimizing representations for individuation, in line with previous findings of an involvement of parietal cortex in face and object individuation tasks^28^.

Prior research has implicated a subset of brain regions within the fronto-parietal network, the so-called “multiple-demand network” in cognitive flexibility^26,29^. It is possible that task-specific optimization across all tasks only takes place within this smaller network of brain areas. We thus conducted an exploratory analysis focusing only on parcels encompassing the core multiple-demand network on the basis of the high-resolution Human Connectome Project atlas^30^ (see Supplementary table 2 for the specific anatomical atlas parcels). Here, similar to frontal cortex overall, we find statistically significant representational optimization only for between-category similarities in the conceptualization task (Fig. S2; repetition 1 vs. 6, between: t(272)=-2.448, p=0.0450, partial eta squared = 0.021557; all other p>0.9, multiple-comparisons corrected for 3 tasks, Individuation: partial eta squared = 0.000651, Categorization: partial eta squared = 0.002086). Taken together, given that both frontal and parietal cortex do not exhibit clear evidence for differential on-task optimization for all three tasks in this study, our results indicate that the neural representations in the fronto-parietal network are not flexibly adjusted according to task demands but contribute only to specific tasks.

Next, we ask whether the medial temporal lobe exhibits flexible transformations towards task-tailored geometries, as it hosts highly pattern-separated, orthogonal representations^18,20,31^ as well as conceptual^32–34^ or abstract representations^35,36^, showing both differentiation and integration of prior knowledge^37^. Here, we find an interaction of task and repetition for between category similarities (F(10, 510)=3.2224, p=0.0004, partial eta squared = 0.0594). However, in contrast to frontal cortex, similarities decrease in the conceptualization task (Fig. 3D; repetition 1 vs. 6, t(510)=2.504, p=0.0378, partial eta squared = 0.012145; linear trend, t(510)=-2.422, p=0.0473 and quartic trend, t(510)=5.105, p=<0.0001). No optimization over repetitions is evident for between category similarities in the individuation task (repetition 1 vs. 6, t(510)=1.500, p=0.1341, partial eta squared = 0.004392), while there is an increase in the categorization task (repetition 1 vs. 6, t(510)=-2.483, p=0.0378, partial eta squared = 0.011944; multiple-comparisons corrected for 3 tasks and 4 trends, respectively). Thus, the medial temporal lobe appears to be involved in increasing differentiation (between categories) in the conceptual task and decreasing differentiation in the categorization task, which seems to run counter the optimal representational structure for both tasks (see below). Therefore, although MTL exhibits transformations of representations through repetitions, based on the available data, the changes do not align with task-tailored, optimal representations.

Lastly, we investigate whether sensory cortices exhibit flexible representational transformations across the three tasks. Sensory representations are often considered stable in order to reliably encode the external world^38^. Hence, finding task-dependent representational changes here would be surprising. Starting with occipital cortex, we indeed find no statistically significant signatures of transformation with repetition towards task-optimal geometries (no significant task × repetition interaction, between category similarities: F(10, 731)=1.7896, p=0.05885, partial eta squared = 0.0239; within category similarities: F(10,1534)=1.0430, p=0.4043, partial eta squared = 0.0068). Occipital cortex does however show a repetition effect independent of the task (main effect of repetition, between: F(5, 731) = 7.8318, p < 0.0001, partial eta squared = 0.0508; within-category: F(5, 1534) = 7.1559, p < 0.0001, partial eta squared = 0.0228; Fig. S3). Between category similarities vary in fashion captured by up to quartic polynomials (t(731)=2.339, p=0.0196, multiple-comparisons corrected for 4 trends), and within category similarities up to cubic polynomials (t(1534)=3.124, p=0.0054, multiple-comparisons corrected for 4 trends), paradoxically increasing on the last repetition (repetition 1 vs. 6, between: t(731)=-4.292, p<0.0001, partial eta squared = 0.024581; within: t(1534)=-3.715, p=0.0002, partial eta squared = 0.008917). While this suggests adjustment of representations, it does not speak to an involvement of occipital cortex in on-task optimization of representational spaces in a task-dependent manner that is optimal to solve the multiple tasks.

We then turned to ventral temporal cortex, which contains sensory representations most selective for the stimuli used here. Here, we find clear signatures of full task-dependent representational reconfiguration predicted by our hypothesis. This was evidenced by highly significant interactions of task and repetition for both between (Fig. 3E; F(10, 2244) = 4.8322, p < 0.0001, partial eta squared = 0.0211; within-category similarity: F(10, 3546) = 7.1779, p < 0.0001, partial eta squared = 0.0198): between category similarities decrease over repetitions in the individuation task (repetition 1 vs. 6, t(2244)=3.167, p=0.0047, partial eta squared = 0.005796; linear trend, t(2244)=-3.691, p=0.0009), while they increase in the conceptualization task (repetition 1 vs. 6, t(2244)=-2.770, p=0.0113, partial eta squared = 0.003408; linear trend, t(2244)=2.647, p=0.0245; quartic trend, t(2244)=4.035, p=0.0002); we find no significant changes in the categorization task (repetition 1 vs. 6, t(2244)=-1.752, p=0.0799, partial eta squared = 0.001366, multiple-comparisons corrected for 3 tasks and 4 trends, respectively). Within category similarities decrease linearly (t(3546)=-5.198, p<0.0001) between repetition 1 and 6 in the individuation task (repetition 1 vs. 6, t(3546)=4.65, p<0.0001, partial eta squared = 0.006061). In contrast, they increase in the categorization (repetition 1 vs. 6, t(3546)=-3.65, p=0.0002, partial eta squared = 0.003743; cubic trend, t(3546)=3.629, p=0.0012) and conceptualization tasks (repetition 1 vs. 6, t(3546)=-5.11, p<0.0001, partial eta squared = 0.007310, linear trend: t(3546)=2.415, p=0.0473, cubic trend: t(3546)=5.504, p<0.0001, multiple-comparisons corrected for 3 tasks and 4 trends, respectively). This suggests that representations in ventral temporal cortex are adjusted over repetitions in a task-dependent way: between and within category similarities are reduced in the individuation task, supporting an improvement in the separability of individual stimuli. In contrast, in the conceptualization task, where visually dissimilar stimuli need to be joined into one concept, between category similarities increase; the same occurs for within category similarities, hence effectively merging face and house representational spaces. In the categorization condition, only within category similarities increase, thus increasing cohesion within categories. These results did not depend on specific ventral temporal cortex sub-regions (no parcel x repetition x task interaction, within category similarities F(60, 4442) = 0.7127, p = 0.954015, partial eta squared = 0.0095; between-category similarity: F(60, 2142) = 0.9090, p = 0.673171, partial eta squared = 0.0248). Further, to control for greater electrode coverage in ventral temporal cortex, we repeated the analysis using subsampling of electrode counts matched to the other brain areas and obtained the same pattern of results (see Supplementary Section 3). Thus, flexibility for the three tasks emerges through gradual optimization towards task-tailored representations in ventral temporal cortex.

Taken together (see Table 1 for an overview, Fig. S4), ventral temporal cortex emerges as the key region supporting all three tasks by flexibly optimizing its representational geometry online to task demands. In the individuation task, parietal cortex additionally adjusts its representation in a task-specific way, by decreasing within and between category similarities over repetitions; occipital and frontal cortex also show task-dependent adjustments of their representations for this task, but no statistically significant signatures of gradual online optimization with repetitions. In the conceptualization task, frontal cortex exhibits an increase in within and between category similarities, supporting joining of visually distinct stimuli into a single concept. The medial temporal lobe instead shows the opposite pattern, decreasing the similarity between faces and houses. Overall, our results suggest that task-tailored representations emerge through gradual optimization over repetitions. The flexibility to carry out multiple tasks on the same set of stimuli thus arises from context-dependent, multipurpose representations in the ventral temporal cortex and is accompanied by task-specific representational optimizations in parietal and frontal cortex.

**Table 1:**
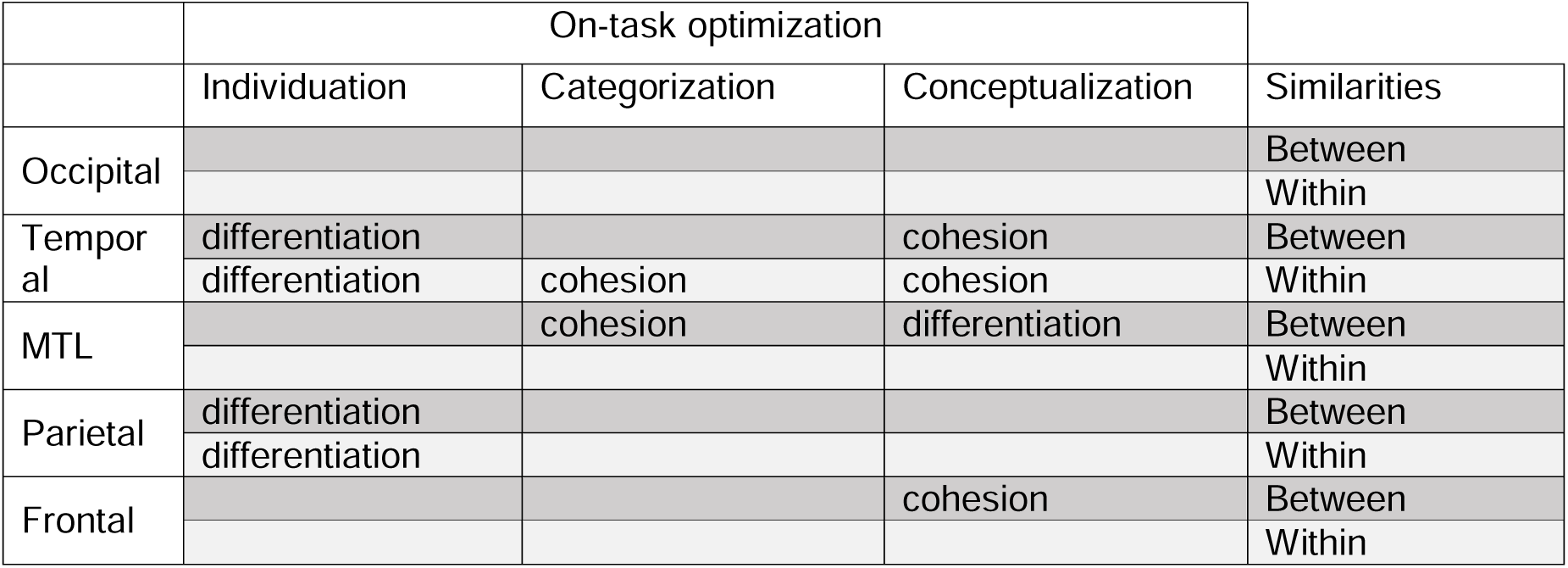
Summary of representational adjustments. “On-task optimization” refers to statistically significant interactions of task and repetition. “Between” refers to between category similarities; “within” refers to within category similarities. “Differentiation” describes decreases in temporal pattern similarities; “cohesion” describes increases in temporal pattern similarities.

|  | On-task optimization |  |  |  |
| --- | --- | --- | --- | --- |
|  | Individuation | Categorization | Conceptualization | Similarities |
| Occipital |  |  |  | Between |
|  |  |  |  | Within |
| Temporal | differentiation |  | cohesion | Between |
|  | differentiation | cohesion | cohesion | Within |
| MTL |  | cohesion | differentiation | Between |
|  |  |  |  | Within |
| Parietal | differentiation |  |  | Between |
|  | differentiation |  |  | Within |
| Frontal |  |  | cohesion | Between |
|  |  |  |  | Within |

### Flexible representational changes are relevant for behavior

Our results suggest that representations are not task-optimal from the outset of a trial, but that task-reconfiguration triggered by the cue requires further task-specific gradual changes through repetitions to optimize representations. What remains unknown is whether the representational optimization we observe is indeed useful and relevant for behavior. We hence pursued two complementary approaches to test whether representational optimization supports higher task accuracy: i) we examined whether the degree of representational optimization across repetitions was greater in subjects with higher behavioral accuracy (between-subjects analysis); and (ii) we tested whether representational optimization predicted trial-by-trial errors (within-subject analysis). We focused on the ad hoc conceptualization task, which provided a sufficient number of participants (n=5) and error trials (≥3 per subject) to enable robust comparisons for the within-subject analysis.

In the conceptualization task, participants must merge two visually distinct object categories, a demand reflected by increased between-category similarity in temporal and frontal cortices. If such optimization of representational geometry contributes directly to behavior, then its strength should scale with accuracy. Specifically, we hypothesized that subjects who were overall better in performing the task (i.e., with accuracy above the group median) would show more pronounced representational optimization across repetitions than subjects with worse performance (i.e., with accuracy below the group median). Consistent with this prediction, we observed a group × repetition interaction in both temporal cortex (F(5,650)=2.2973, p=0.0438) and frontal cortex (F(5,585)=4.0928, p=0.0012). In both regions, optimization was significantly stronger in the high-accuracy group compared with the low-accuracy group (ventral temporal cortex: t(650)=2.954, p=0.0032; frontal cortex: t(585)=3.196, p=0.0015). No other area showed statistical significance for such an effect (all p>0.2288), highlighting that behaviorally relevant representational changes are localized to temporal and frontal regions.

As a complementary approach, we compared representational optimization between correct and incorrect trials. In temporal cortex, error trials showed a decrease in between-category similarity over repetitions, in contrast to correct trials (accuracy × repetition interaction: F(5,573)=4.8520, p=0.0002; repetition 1 vs. 6: t(573)=2.279, p=0.0460). A similar accuracy × repetition interaction was found in frontal cortex (F(5,532.26)=2.3462, p=0.0401), driven by a reduction in between-category similarity on error trials (repetition 1 vs. 6: t(532)=-2.091, p=0.0370). The medial temporal lobe also exhibited an accuracy × repetition interaction (F(5,118)=2.7038, p=0.0237), although the effect did not reach significance when comparing repetitions 1 and 6 directly (t(118)=-0.446, p=0.6564).

Finally, we examined within-category similarities, where only temporal cortex had shown evidence of representational cohesion in the conceptual task. Again, optimization was behaviorally relevant, as evidenced by a group × repetition interaction (F(5,1445.05)=3.3508, p=0.0052), with stronger optimization in high-accuracy than low-accuracy participants (t(1445)=2.689, p=0.0073). No other area showed comparable, statistically significant effects (all p>0.0665).

To assess whether behavioral changes simply occurred through reductions in the amplitude of evoked responses, we carried out a supplementary analysis focusing on repetition suppression. We find that although repetition suppression occurs in some areas, ventral temporal cortex does not show consistent reduction or changes in amplitude over repetitions for all the three tasks (see Supplementary Section 4, Supplementary Figures S7 and S8). Furthermore, behavior does not correlate significantly with response amplitudes across repetitions in any area (see Supplementary Section 4). This indicates that gradual representational optimization in the ventral temporal cortex is a key for behavioral performance, which cannot be explained solely by amplitude changes.

Overall, our findings suggest that successful behavior in the ad hoc conceptualization task depends on the appropriate cohesion of representations both within and between categories. Failures to optimize representational geometries across repetitions - manifesting as insufficient alignment of the two types of representations that must be combined in this task - lead to behavioral errors and reduced accuracy.

## Discussion

Taken together, our results reveal that the ability to flexibly perform multiple tasks is supported by task-tailored representational geometries that, instead of a one-shot transformation, emerge through gradual optimization over repeated experience. Remarkably, the only brain area that showed evidence of flexible adjustments for all tasks studied here was the ventral temporal cortex. Hence, sensory cortices do not simply represent the external world, but exhibit context-dependent flexible optimization on a trial-by-trial basis. In addition to the task-dependent changes in ventral temporal cortex, representational transformations are also evident in larger distributed networks, including frontal cortex, that configure themselves in task-specific ways. Overall, this indicates that mental flexibility, a hallmark of human intelligence, is instantiated by reconfiguring sensory neuronal representations for multiple tasks.

The flexibility of ventral temporal cortex revealed by our data departs from the traditional view of this region as a stable perceptual frontend whose representations are merely read out by downstream systems. Much of that view stems from studies relying on passive viewing and/or anesthetized animals^39–42^, where task demands cannot reshape sensory codes. In contrast, emerging evidence^43^ ^44–46^ - and our results - show that sensory cortex can undergo rapid, context-dependent plasticity, adjusting representational distances in ways that improve task performance. Such adjustments in sensory cortices are computationally advantageous: changes at this stage can exploit fine-grained feature dimensions unavailable to later regions^47^ and can be deployed already early during processing^46^. Thus, ventral temporal cortex should be understood not only as a hub for high-level visual representations, but also as an active contributor to goal-dependent optimization of representational geometry.

The available evidence suggests that frontal and parietal cortex adjusted their representations for only a subset of tasks - but not all three. Thus, we found no indication in our data that these regions carry out the kind of general, rapid representational reconfiguration observed in ventral temporal cortex. Our results do not necessarily contradict evidence that fronto-parietal areas, in particular the multiple demand network^27^, encode multiple task sets; rather, our results suggest that changing representations in these areas may require more prolonged experience (as in perceptual^48–50^ or category learning^51–53^) than what was probed here. Alternatively, or additionally, fronto-parietal areas may be more involved in carrying out multiple tasks if they span multiple modalities and/or require very distinct sensorimotor mappings. Under rapid task switching, these regions may rely on representations shaped earlier in the processing hierarchy instead of restructuring their own geometry on a trial-by-trial basis.

Flexible representational adjustments in ventral temporal cortex were accompanied by task-specific refinements in the fronto-parietal cortex, in line with previous findings that show task specialization in these areas^54–56^. Frontal cortex was particularly involved in the formation of an abstract, ad hoc concept/category by increasing representational cohesion between the categories that needed to be merged. It is possible that frontal cortex supports ventral temporal cortex in this task that may exceed what is possible in terms of conjoining visually very distinct object categories in high-level visual cortex. Parietal cortex showed on-task representational adjustments by increasing differentiation of representations in the individuation task. Parietal cortex has previously been shown to be involved in the individuation of complex objects, including faces^28^. Yet, parietal cortex is of course also implicated in the formation of categories^51^, a task for which we did not find on-task optimization. It is possible that this discrepancy again relates to long-term acquisition of categories which often involves extensive training, while our tasks were based on trial-by-trial instruction and required flexible switching. Beyond these stimulus-evoked adjustments of representational geometry, parietal cortex also showed task-dependent pre-stimulus high-gamma band activity (F(2,293)=4.36, p=0.0136), with greater amplitude during ad hoc conceptualization than individuation (t(293)=−2.87, p=0.013). This anticipatory modulation may reflect attentional engagement or maintenance of the currently relevant task set.

MTL regions also showed task-dependent changes in representational geometry, but these adjustments followed a pattern opposite to the one observed in cortex: between-category similarities increased in the categorization task (where differentiation would have been optimal) and decreased in the ad hoc conceptualization task (where cohesion would have been optimal). Such inversions are consistent with competitive interactions between MTL and cortex, as previously reported^37^, and parallel accounts from the non-monotonic plasticity framework, where highly similar representations become decorrelated most strongly in MTL^57^. An alternative interpretation is that representational changes in the MTL may reflect functions other than immediate task performance, such as maintaining pattern stability to support memory, which has been linked to reduced representational change across repetitions^58^. Finally, hippocampal adjustments may simply operate on slower timescales than the rapid cortical dynamics captured here, with more pronounced task-appropriate changes emerging only after a larger number of repetitions than were feasible in this paradigm^59^. Further studies will be necessary to pin down the contribution of MTL to flexible multi-task performance.

Theoretical work on multitask learning suggests that reusing shared representations supports rapid generalization and compositional behavior^4,5^, but that these benefits come with costs: shared representations can interfere with task-specific processing and reduce efficiency^8,60^. This offers a natural explanation for the dissociation we observe between rapid behavioral multitasking and the lack of evidence for *immediate* task-tailored geometries. One- or even zero-shot task performance may rely on the rapid readout of a rich, high-dimensional representational space, possibly by the fronto-parietal network. This may allow for high (but not perfect) accuracy without requiring instantaneous reconfiguration. Transforming these shared representations into task-specific geometries instead became detectable in our data only over the course of a few stimulus repetitions, possibly to avoid interference. Consistent with this view, neural networks develop task-tailored (“rich”) representations only later in training^61^, with gradual adjustments providing robustness to noise and preventing idiosyncratic inputs from destabilizing existing structure^12^. Models that succeed on rapid timescales instead often rely on task-agnostic, high-dimensional representations^62–64^. A switch of strategies between the first and later repetitions would essentially balance the computational benefits of sharing representations and separating tasks.

Our analyses revealed that temporal codes are a powerful dimension of neural representation in human cortex, allowing sensory representations to flexibly transform according to different task demands. In addition, a spatial population analysis (see Supplementary Section 2) revealed a complementary, contrastive code^65,66^, where representations became progressively differentiated during individuation in ventral temporal and parietal cortex, whereas we found no statistically significant evidence for comparable optimization in the other tasks. Such a contrastive code is thought to instantiate a single objective function whereby representational separation between different stimuli is enhanced^67,68^. This can improve discrimination, robustness to noise, and downstream readout, making this code efficient for a specific objective such as individuation. However, such specialization may reduce flexibility for situations requiring the same stimuli to be grouped or related differently, especially when tasks are interleaved. Temporal patterns in ventral temporal cortex, by contrast, became tailored to all three tasks under study. It is possible that temporal multiplexing^19,69–71^ expands the dimensionality of representations, allowing the same neural population to support multiple, potentially conflicting transformations within a shared representational space.

However, these mechanisms need not be confined to representational changes within individual cortical areas. Rapid task switching could additionally involve task-dependent attentional shifts across regions, preferentially weighting representations whose existing geometry is specialized or well suited to the current goal^72^. Although our data do not directly demonstrate attentional shifts, the task-specific pattern of effects observed in parietal cortex in individuation and in frontal cortex in ad hoc concept formation, alongside flexible optimization across all three tasks in ventral temporal cortex, is consistent with this possibility.

In conclusion, our results show how the brain deals with the challenges of multiple tasks. A high degree of representational flexibility enables balancing multiple and often conflicting demands. The plasticity we find even on short timescales reaches further down the processing hierarchy than previously thought, i.e., all the way to visual cortex. Hence, our work suggests that neural codes in the sensory cortex are not stable or functionally specialized, but instead can be used to support flexible computations and generalization^73,74^. Ultimately, the widespread and profound changes in response to task demands may explain how we can rapidly change our mind when our goals change – and thus, mental flexibility, a hallmark of intelligent behavior.

## Material and Methods

### Participants

Intracranial electrophysiological recordings were obtained from 14 patients (6 female, 3 left-handed, age: average 33.35 years, range 21-60) undergoing invasive monitoring of pharmacologically resistant epilepsy. Sex was obtained from participants’ clinical records; gender identity was not collected. Testing took place at North Shore Long Island Jewish University Hospital (LIJ), Manhasset, NY, USA (n=9), New York University (NYU) Langone Medical Center, NY, USA (n=2) and the Epilepsy Department of the Grenoble Alpes Hospital, Grenoble, France (n=3). Each participant provided informed consent before the experiment, following the guidelines of the Declaration of Helsinki. Participants were informed that participating in the study would not affect their clinical treatment and that they could withdraw at any time without affecting their medical care. All procedures were approved by the Institutional Review Boards at the Feinstein Institute for Medical Research, New York University Langone Medical Center, and Grenoble Hospital, and by the National French Science Ethical Committee.

### iEEG data acquisition

All electrode implantations, including the decision to implant and the location of the implantation were solely determined based on clinical grounds. Platinum iridium macroelectrodes (Ad-Tech Medical Instrument at LIJ, NYU centers and Dixi Medical at Grenoble) were implanted sub-durally with a 2-3-mm-diameter exposed surface. Macroelectrodes were arranged as grid arrays (6 x 8, 8 × 8 contacts, 10- or 5-mm center-to-center spacing), linear strips (1 × 4/6/8/12 contacts), or depth electrodes (1 × 8/10/12 contacts), or some combination. Subjects at LIJ, NYU had surface grid, strip and depth electrodes, and subjects at Grenoble had only depth electrodes implanted. At LIJ, intracranial data were collected using a Brainbox EEG-1166 system (Braintronics) at a sampling rate of 1000 Hz. These signals were referenced online to a subgaleal electrode at the vertex. At NYU, neural data were collected using a Nicolet ONE amplifier (Natus) at a sampling rate of 512 Hz and referenced online to a skull-mounted screw. At Grenoble Hospital, data were collected using a Micromed system at a sampling rate of 512 Hz using a reference electrode located in the white matter. All data were continuously stored along with stimulus and timing markers to enable offline synchronization. Channels were excluded based on visual inspection if they exhibited interictal spikes, electromagnetic interference, or other noticeable signal artifacts; if they were located within the clinically defined seizure onset zone; or if reliable anatomical localization in the MRI was not possible (3% of the electrodes).

### Stimuli and Experimental Procedures

#### Localizer

The stimuli were presented using Presentation (Neurobehavioral Systems) on a laptop at bedside. Subjects responded with their dominant hand, using a keyboard. In the visual localizer task, participants were exposed to images from various visual categories in a random sequence while maintaining fixation on a central cross. To ensure attention, they performed a cover task that varied slightly across sites. At LIJ and NYU, participants were asked to respond whenever an identical image was presented twice consecutively. Stimuli belonged to 6 different categories: Face, House, Object, Scrambled pattern, Body, and Eye. Each image appeared for 250 ms with a 750 ms inter-stimulus interval (ISI). At Grenoble Neurological Hospital, participants were asked to respond whenever an image of a fruit was shown. Stimuli included Face, House, Tools, Scrambled pattern, Animal, Scene, Pseudowords, and Consonants, each presented for 200 ms with a 750 ms ISI. Participants and investigators were not blinded to experimental conditions. The categories pseudowords, scene, eye, and animal were not used for further analyses.

#### Main Experiment

For the main experiment, 284 natural images (71 exemplars each from faces, houses, objects, animals) were normalized to have identical Fourier amplitude spectra. Degraded versions of natural images were created by blending their Fourier phase spectra with random phase spectra, separately for each RGB color channel. Contextual cues were shown without any phase scrambling, but all repetitions were presented with 50% phase scrambling. The stimuli were presented using Presentation (Neurobehavioral Systems) on a laptop at bedside. Subjects responded with their dominant hand, using a keyboard.

On each trial, subjects were exposed to a series of 6 images (faces/houses) presented in a sequence one after the other, and were asked to identify whether a deviant was presented among the repetitions. The subjects performed three tasks and the correct response to identify a deviant depended on the task. The contextual cue which instructed the task demand on a trial-by-trial basis was followed by repeated stimuli of two different visual categories: faces and houses. The three tasks were as follows: 1) individuation - where the repeating stimuli are a specific face/house identity and the subjects report any different identity as a deviant, 2) categorization - where the repeating stimuli are from within a visual category (any face/house) and the subjects report any difference in a visual category as a deviant, and 3) conceptualization - where the repeating stimuli are either faces or houses and here any non-face or non-house object is considered a deviant. The different tasks as well as the cues used for each task were explained both verbally and in written form before the experiment.

During the study, each trial began with the contextual cue (1000 ms) indicating the task to be performed in the subsequent images, followed by a 2000 ms inter-stimulus interval (ISI). Six images were then presented sequentially (200 ms each, ISI = 1000–1300 ms), after which participants indicated whether a deviant had occurred. After the subject responded, a 1500– 2000 ms inter-trial interval followed before the presentation of the next cue. Participants and investigators were not blinded to experimental conditions, as task conditions were explicitly cued.

Each subject completed 3-4 blocks of 35 trials each, except subject grenoble_suj2 who was presented only one block and grenoble_suj3 who was presented with two blocks. Each of the three tasks, individuation (face/house), categorization (face/house), conceptualization had an equal number of trials within a block (7 trials per condition per block). 28.75% of the trials (10 out of 35) contained a deviant. Unless otherwise specified, electrophysiological analyses were restricted to trial sequences without deviants were subjects responded correctly that no deviant was presented

### Quantification and Statistical Analyses

#### Anatomical analysis: electrode localization and regions of interest

Localization of electrodes was done using using the iELVIs toolbox^75^. Pre-surgical T1-weighted, anatomical MRIs were reconstructed and segmented using FreeSurfer (version 5.3)^76^. The postsurgical high density thin-slice computed tomography (CT) images or post-surgical anatomical MRIs were co-registered to the pre-surgical MRI. Electrodes were identified in CT scans using Bioimagesuite^77^ and projected on to the pial surface. Sub-dural electrodes were corrected for post-operative brain-shift using the Dykstra method^78^. Brain-shift correction was not done for depth electrodes^75^. Following this, electrode locations were mapped on to the Desikan-Killiany atlas^79^ to obtain parcel names for each electrode. Owing to the variable coverage of iEEG data, parcels were combined to obtain coarse clustering of brain regions (as in previous studies^80,81^) for further analyses (Occipital cortex, Ventral temporal cortex, Parietal cortex, Medial temporal lobe, Frontal cortex). The parcels per cluster were selected based on Desikan-Killiany Tourville^82^. The boundaries of the ventral temporal cortex were chosen based on previous work^83^ (see Supplementary Table 1, Supplementary Fig. 1). For a control analyses, we also mapped the sub-dural electrodes onto the surface-based Human Connectome Project (HCP)^30^ atlas. The parcels defined as the core multiple-demand network were combined into one region of interest^29^ (see Supplementary Table 2 for parcels in this ROI). Further, depth and subdural electrodes were combined for all cluster-level analyses which is standard practice in several human intracranial electrophysiology studies^84–86^. The numbers of implanted electrodes from each electrode type are reported separately in Supplementary Table 3.

#### Behavioral analysis

Performance (accuracy and RT) on different tasks was calculated per subject and task after outlier detection. Outliers in the RT were identified at the single-trial level using the Croux and Rousseeuw method^87^ with a threshold of 3. Only correct trials were included in the main RT analyses. Task differences in accuracy and RT were assessed with repeated-measures ANOVAs, with task as the within-subject factor. To investigate whether image number in the sequence (repetition) influenced RT for deviance detection, we conducted post-hoc trend analyses following the ANOVA, restricted to correctly detected deviants.

#### Electrophysiology Analysis

##### Preprocessing

All data preprocessing was done in Matlab (2018b, The Mathworks) using the Fieldtrip toolbox (version 20190329)^88^ and was done in line with recommendations in the field^89^. Channels were visually examined and excluded if they contained interictal spikes, electromagnetic artifacts, or if they were in clinically determined seizure onset, or if they could not be reliably localized in the anatomical scan. Line noise was removed using a discrete Fourier transform filter (DFT filter) where the powerline component’s amplitude is estimated by fitting a sine and cosine at the specified frequency and the signal is subtracted from the data. For the data collected in the USA, this procedure was done at the default power line value of 60 Hz and its harmonics (120 and 180 Hz). For the data collected in Europe, 50 Hz and its harmonics (100, 150, 200 Hz) were used as the specified frequency. Each subject’s data was demeaned and re-referenced to the average of all artifact-free electrodes and resampled to 500 Hz after an anti-aliasing low pass filter was applied^88^.

##### High-Gamma Band Amplitude

For the main analysis, we focused on high-gamma band (HGB) activity, which closely correlates with neuronal spiking and population firing rate ^16,17,90^. Time–frequency decomposition was performed in the 70–200 Hz range using a multi-taper convolution method with Hanning tapers, sampled in 5 Hz steps. To correct for the 1/f power drop-off, power estimates at each frequency were weighted by the square of their respective frequency values ^91^ and subsequently averaged across frequencies.

HGB Power was baseline corrected on a trial-by-trial basis^92^ using the decibel (dB) conversion method^93^. Specifically, for each trial, power values were normalized by the mean activity of a designated baseline period and then log-transformed (10*log10). For the main dataset, the baseline period was defined as –300 ms to –50 ms relative to cue onset. This window was selected to ensure that normalization was constant across repetitions and unaffected by task-context information. For the localizer dataset, baseline was defined as – 300 ms to –50 ms before each image presentation^40,41,89^. For the supplementary analysis, mean HGB amplitude was computed by averaging across trials, frequencies, and the first 250 ms of the electrode’s onset response. This measure was calculated separately for each electrode and task. Unless otherwise specified, all analyses were restricted to trials without deviants and to trials with correct behavioral responses.

##### Identification of visually responsive sites and onset latency

The localizer dataset was used to identify visually responsive sites. For each electrode and category, we compared post-stimulus HGB power (0–500 ms) with baseline activity (–300 to –50 ms) using a two-sided Wilcoxon rank-sum test on every 5 ms time bin (α = 0.01). To correct for multiple comparisons across electrodes and time points, we applied the False Discovery Rate^94^ with a q-value threshold of = 0.01^93^. We then defined a Response Index (RI) as the number of visual object categories that elicited a significant average HGB response at each electrode. Electrodes were classified as visually responsive if they showed significant responses to at least one category (i.e., RI>0) for 10 or more consecutive time bins (i.e., ≥ 50 ms of activity above baseline)^95^.

##### Within and Between Category Similarities

To quantify representational similarity, we analyzed single-trial HGB amplitude from 0–1000 ms following image onset. Data were downsampled using a running mean over 20 ms windows. Following approaches from the hippocampal pattern separation literature^18,19^ temporal similarity was defined as the Pearson correlation (over time) between neural responses elicited by different images.

For within-category similarity, correlations were computed separately for each task, visual category (faces, houses), and repetition. This was performed for each visually responsive electrode in each subject. However, to test whether this selection affected our conclusions, we additionally included visual responsiveness as a factor and analyzed all electrodes; the main pattern was preserved, although task-tailored changes were clearest in visually responsive electrodes (Supplementary Section 5). Pairwise correlations between images of the same category were averaged to yield within-category similarity values for faces and houses separately. These values were then averaged to obtain a single within-category similarity estimate per task and repetition for each subject. For between-category similarity, correlations were computed between responses to each pair of face and house images presented to a subject, again separated by task and repetition. We validated the reliability of this measure using split-half analyses of an independent localizer dataset, which showed comparable reliability across regions, except in occipital cortex (Supplementary Section 6, Fig. S9). Pairwise correlations were averaged within each subject and electrode to yield a between-category similarity measure. All analyses were restricted to trials with correct behavioral responses, except when explicitly relating neural similarity to behavior.

##### Relationship with behavior

To test whether failures of representational optimization lead to errors in task performance, we conducted two complementary analyses: (a) a between-subjects comparison of representational changes in participants with higher versus lower accuracy, and (b) a within-subject comparison of correct versus error trials. Analyses were restricted to the conceptual task, as this condition provided a sufficient number of subjects (n = 5) and error trials (≥ 3 per subject) to permit reliable statistical testing. The minimum trial requirement was based on prior electrophysiological studies^96^.

For the between-subjects analysis, all trials in which stimuli were repeated without a deviant (including both correct and incorrect responses) were used to calculate amplitude as well as within- and between-category similarity. Subjects were then divided into two groups (high vs. low accuracy) based on median task accuracy, calculated as percent correct across all repeated and deviant trials.

For the within-subject analysis, error trials were defined as repetitions without a deviant in which the subject incorrectly reported a deviant (“false alarms”). Within- and between-category similarities were calculated for these error trials, following the procedure described in the previous section. To compare against correct trials, we subsampled correct repetitions to match the number of error trials for each subject, ensuring balanced trial counts^95^.

#### Statistical Analyses

Statistical analyses were done in R (4.2.1) and Matlab (v2018a, 2018b). To determine whether and where the representations are task-tailored at repetition 1, we first fitted a linear mixed effects models to within and between-category similarities, using the lme4 package (version 1.1.35.1)^97^. We predicted similarity at repetition 1 with the factors ‘task’ (3 levels, individuation/categorization/conceptualization) and region cluster (5 levels, frontal, parietal, occipital, ventral temporal cortex, medial temporal lobe) as and their interaction as fixed effects, ‘subjects’ as a random effect, and ‘electrodes’ as nested random effect to the average within and between category similarities, respectively. This hierarchical structure of the statistical model captures the structure of the data, accounting for differential variability between subjects and within electrodes, and allows for generalization to the population^98,99^. We identified outliers on the basis of normalized residuals exceeding a threshold of 4.5 and removed the respective electrode from all conditions before re-fitting the model. Subsequently, we fitted one model per region of interest. We then tested the contrasts that define task-tailored representations: Categorization > Individuation and Conceptualization > Individuation for within-category similarity and Conceptualization > Categorization and Conceptualization > Individuation for between-category similarity for each region. After that, we combined the p-values using a combination test for within and between-category similarity, respectively. This was done using the Stouffer method implementation in the poolR package^100^. Because the representational geometry for the tasks we used is dependent on both within and between category together to be optimal and task-tailored, we further combined the p-values from both within and between similarities using the Stouffer method in a combination test. This tests whether the contrasts Categorization > Individuation and Conceptualization > Individuation for within-category similarity and Conceptualization > Categorization and Conceptualization > Individuation for between-category similarity is significant together. As a complementary analysis, we also computed mean difference between Categorization - Individuation and Conceptualization - Individuation for within-category similarity and the mean difference between Conceptualization - Categorization and Conceptualization - Individuation for between-category similarity. Lastly, to ask whether both the within and between category similarity differences are significantly above 0 (top-right quadrant in Fig. 2B), we computed one-sided t-tests.

To determine how task and repetition affected within and between category similarities, we fitted linear mixed effects models to the average within and between category similarities, respectively. We predicted similarity and amplitude with the factors ‘task’ (3 levels, individuation/categorization/conceptualization) and ‘repetition’ (6 levels, repetition 1-6) and their interaction as fixed effects, ‘subjects’ as a random effect, and ‘electrodes’ as nested random effect (intercept), similar to previous studies^101,102^. For within and between similarities, respectively, we first fitted a model that included all regions of interest as an additional factor to determine whether regions differed in terms of how task and repetition affected similarities. We identified outliers on the basis of normalized residuals exceeding a threshold of 4 and removed the respective electrode from all conditions before re-fitting the model. Subsequently, we fitted one model per region of interest. In case a model did not converge, we determined whether a model that included subjects and electrodes as separate (not nested) random effects, or a model that included only subjects or only electrodes, respectively, as random effects, converged and provided a better model fit on the basis of the relative log likelihood between the models (based on the Akaike Information Criterion, AIC)^103^. For each fitted model per region (for amplitudes, within and between similarities), we identified outliers on the basis of normalized residuals exceeding a threshold of 4.5 and removed the respective electrode from all conditions before re-fitting the model. For the analysis with error trials, we fit a linear mixed-effect model with ‘correctness’ (2 levels, correct and incorrect), and ‘repetition’ for only the conceptualization task. This was done for both within and between category similarities. Outlier electrodes detected from the main within and between category similarity analysis were removed from this analysis prior to fitting the model. To obtain *p*-values, we performed type III analyses of variance (ANOVA) of the model terms and their interactions using Wald’s *F*-test and determined degrees of freedom following Kenward and Roger^104^ using the lmerTest package (version 3.1.3)^105^. Posthoc tests and trend analyses (up to 4^th^ order polynomials) were performed on estimated marginal means using the emmeans package^106^ (version 1.10.0) (https://cran.r-project.org/web/packages/emmeans/index.html). All plots show estimated marginal means and model-based standard error of means from the emmeans package. All posthoc tests and trend analyses were corrected for multiple comparisons using Holm’s sequential Bonferroni procedure^107^. All posthoc tests were two-sided comparisons. For pairwise contrasts from the linear mixed-effects models, we report test-statistic-based partial eta squared as effect size, using the model-derived denominator degrees of freedom. Sex was not included as a factor in the statistical analyses because the clinically recruited sample was neither designed nor powered for sex-based comparisons.

## Data availability

Source data are provided with this paper. The data to reproduce the figures in this study have been deposited in the Figshare database (link will be made available after journal acceptance). Further information and requests for resources should be directed and will be fulfilled by Caspar M. Schwiedrzik upon reasonable request.

## Code availability

No custom code was developed specifically for this study. Analyses were performed using standard functions and publicly available software packages as described in the Methods.

## Author Contributions

**T.N.:** Conceptualization; Methodology; Validation; Formal analysis; Investigation; Writing - Original Draft; Writing - Review & Editing; Visualization. **A.F.C-P.:** Formal analysis; Visualization; Writing - Review & Editing; **P.M.:** Investigation; Writing - Review & Editing; **J.R.V.:** Investigation; Writing - Review & Editing; **M.P.B.:** Investigation; Writing - Review & Editing; **P.K.:** Investigation; Writing - Review & Editing; **T.T.:** Investigation; Writing - Review & Editing; **O.D.:** Investigation; Writing - Review & Editing; Funding acquisition; **L.M.:** Conceptualization; Methodology; Validation; Investigation; Writing - Review & Editing; Supervision; Project administration; **C.M.S.:** Conceptualization; Methodology; Validation; Formal analysis; Investigation; Writing - Original Draft; Writing - Review & Editing; Supervision; Project administration; Funding acquisition.

## Acknowledgements

We would like to thank Ashesh D. Mehta, Jean-Philippe Lachaux, and Werner Doyle for contributing their unique expertise to the success of this study. We are grateful to Benjamin Y. Hayden for his insightful feedback and valuable suggestions during the review process of this manuscript. We also thank Rüdiger Ludwig for IT support. This research was supported by the Deutsche Forschungsgemeinschaft (DFG, German Research Foundation) - Project-ID 454648639 - SFB 1528 - Cognition of Interaction, project A04, and the Emmy Noether Program (SCHW1683/2-1) awarded to C.M.S.; P.M. was supported by Swiss National Science Foundation grant 148388; L.M. was supported by the Max Planck Society. This work was also supported by Finding A Cure for Epilepsy and Seizures (FACES; to O.D.). The funders had no role in study design, data collection and interpretation, decision to publish, or preparation of the manuscript.

