## Supplementary Material for "Multiple task-demands flexibly optimize neural geometry in human ventral temporal cortex"

#### Tables

| Cluster Names | Parcel Names | Number of visually responsive electrodes | Number of implanted electrodes | Number of subjects |
|---|---|---|---|---|
| Occipital Cortex | cuneus, lateraloccipital, lingual, pericalcarine | 45 | 64 | 10 |
| Ventral Temporal Cortex | inferiortemporal, middletemporal, parahippocampal, superiortemporal, temporal_fusiform, temporalpole | 140 | 514 | 14 |
| Parietal Cortex | inferiorparietal, postcentral, superiorparietal, supramarginal | 61 | 243 | 8 |
| Medial Temporal Lobe | amygdala, hippocampus, entorhinal | 33 | 96 | 10 |
| Frontal Cortex | caudalmiddlefrontal, frontalpole, lateralorbitofrontal, medialorbitofrontal, parsopercularis, parsorbitalis, parstriangularis, precentral, rostralmiddlefrontal, superiorfrontal | 125 | 436 | 11 |

Supplementary Table 1: Assignment of anatomical parcels into clusters

List of clusters and the respective anatomical parcels from the Desikan-Killiany atlas, the number of visually responsive and implanted electrodes, respectively, across subjects and the number of subjects that contributed to the visually responsive electrodes per cluster.

| Cluster Name | Parcel Names | Number of visually responsive electrodes |
|---|---|---|
| Core Multiple-demand Network | 'a9-46v';'p9-46v' ;'IP1' ;'IP2' ;'IFJp' ;'i6-8' ;'PFm'; 'AVI' ; '8BM';'8C'; | 17 |

Supplementary Table 2: Assignment of anatomical parcels into the core multiple-demand network

List of anatomical parcels from the Human Connectome Project atlas and the number of visually responsive electrodes across subjects.

| Cluster | Depth | Subdural | Total electrodes |
|---|---|---|---|
| Occipital | 14 | 50 | 64 |
| Ventral Temporal Cortex | 214 | 300 | 514 |
| Parietal Cortex | 19 | 224 | 243 |
| Medial Temporal Lobe | 86 | 10 | 96 |
| Frontal Cortex | 39 | 397 | 436 |

Supplementary Table 3: Number of implanted electrodes in each anatomical cluster, separated by electrode type

Electrodes were classified as “depth” for stereotactic depth electrodes and “subdural” for grid or strip electrodes.

### Figures

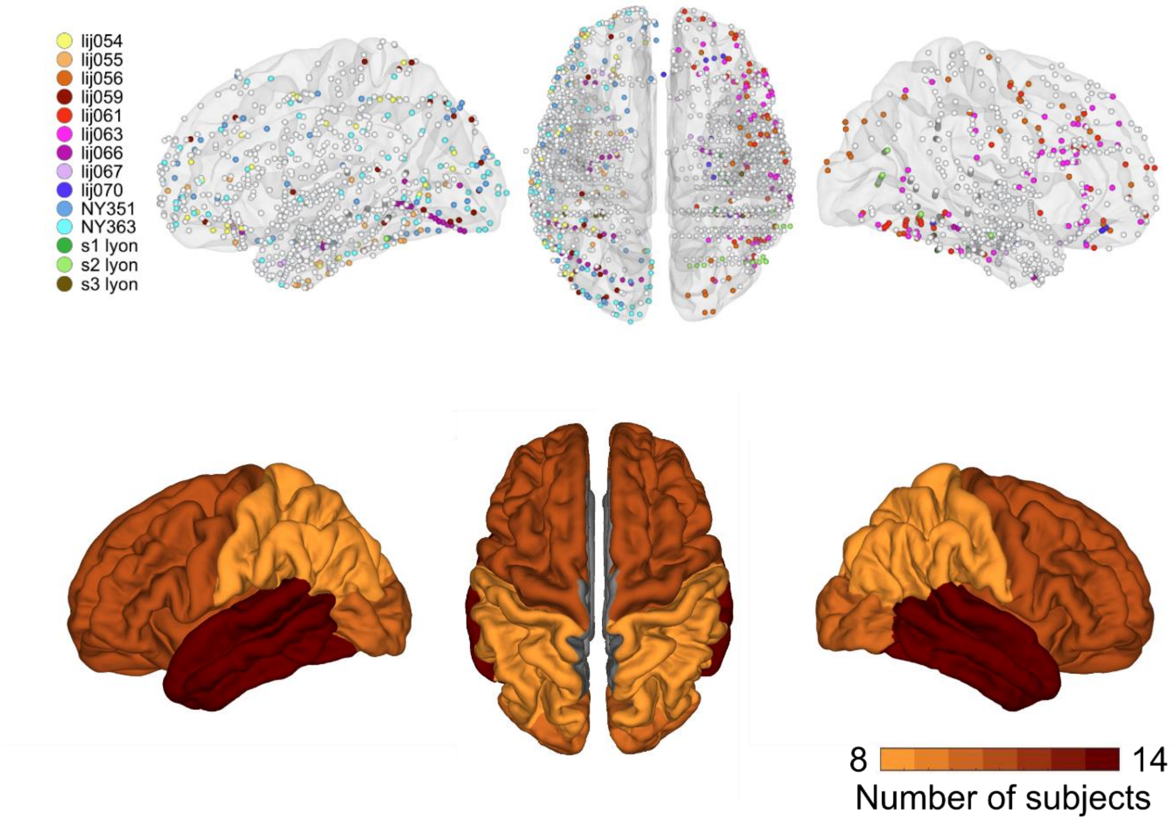


<u>**Fig. S1: Electrode coverage across all subjects**</u>

Upper panel: Electrophysiological recordings were obtained from 2003 electrodes (1957 localized and shown) across 14 epilepsy patients. For analyses, 404 visually responsive electrodes were selected based on an independent localizer. Colored electrodes indicate visually responsive sites (color-coded by subject); gray electrodes represent non-visually responsive ones. Lower Panel: Number of subjects that contributed to each cluster for the main analysis. At least 8 of 14 subjects contributed to each cluster.

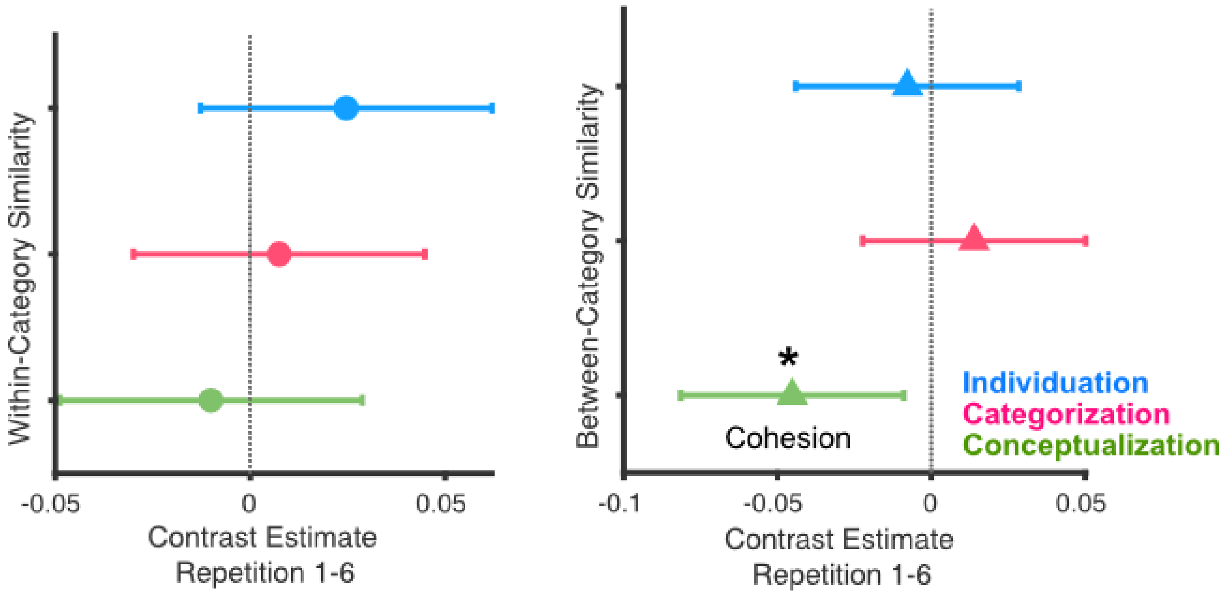


**Fig. S2: Representational optimization for the tasks in the core multiple-demand network**

The core multiple-demand network exhibits increasing cohesion between categories, i.e., between category similarities decreased with repetition in the conceptualization task (repetition 1 vs. 6, between: $t(272)=-2.448$, $p=0.0450$, multiple-comparisons corrected for 3 tasks), while we found no evidence for representational optimization for any of the other tasks (all $p>0.9$, multiple-comparisons corrected for 3 tasks). The x-axis indicates contrast estimates of the post-hoc test between repetition 1 and 6, along with model-based standard error bars. Positive x-values indicate differentiation over repetitions, i.e., decreasing similarity over repetitions, and negative values indicate cohesion over repetitions. Blue indicates individuation task (In), pink indicates categorization task (Ca) and green indicates conceptualization task (Co). The left panel shows representational optimization in terms of within-category similarity and the right in terms of between-category similarity. * $p<0.05$.

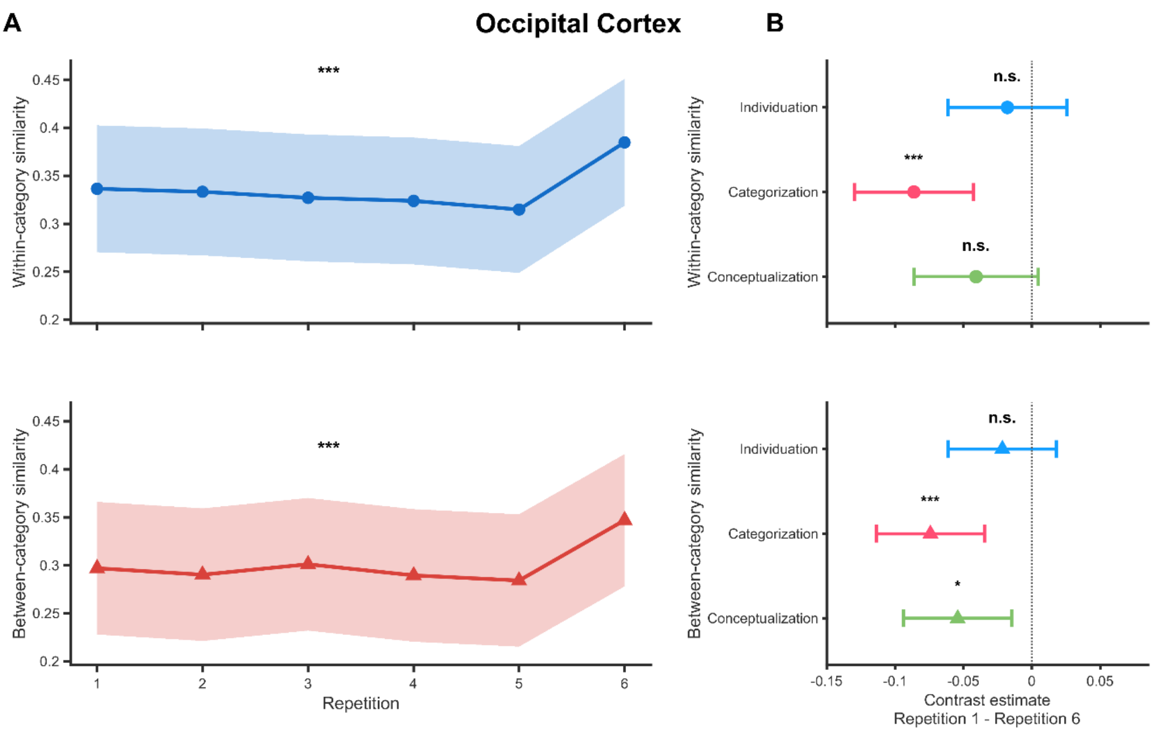


**Fig. S3: Representational similarity changes across repetitions in occipital cortex**

A) Task-averaged estimated marginal means for within- and between-category similarity across repetitions in occipital cortex; shading indicates ±SE. (B) Task-specific repetition 1–6 contrasts; negative estimates indicate increased similarity at repetition 6. Points show contrast estimates and error bars show 95% confidence intervals. P-values were Holm-corrected across tasks (*p < 0.05, ***p < 0.001; n.s., not significant).

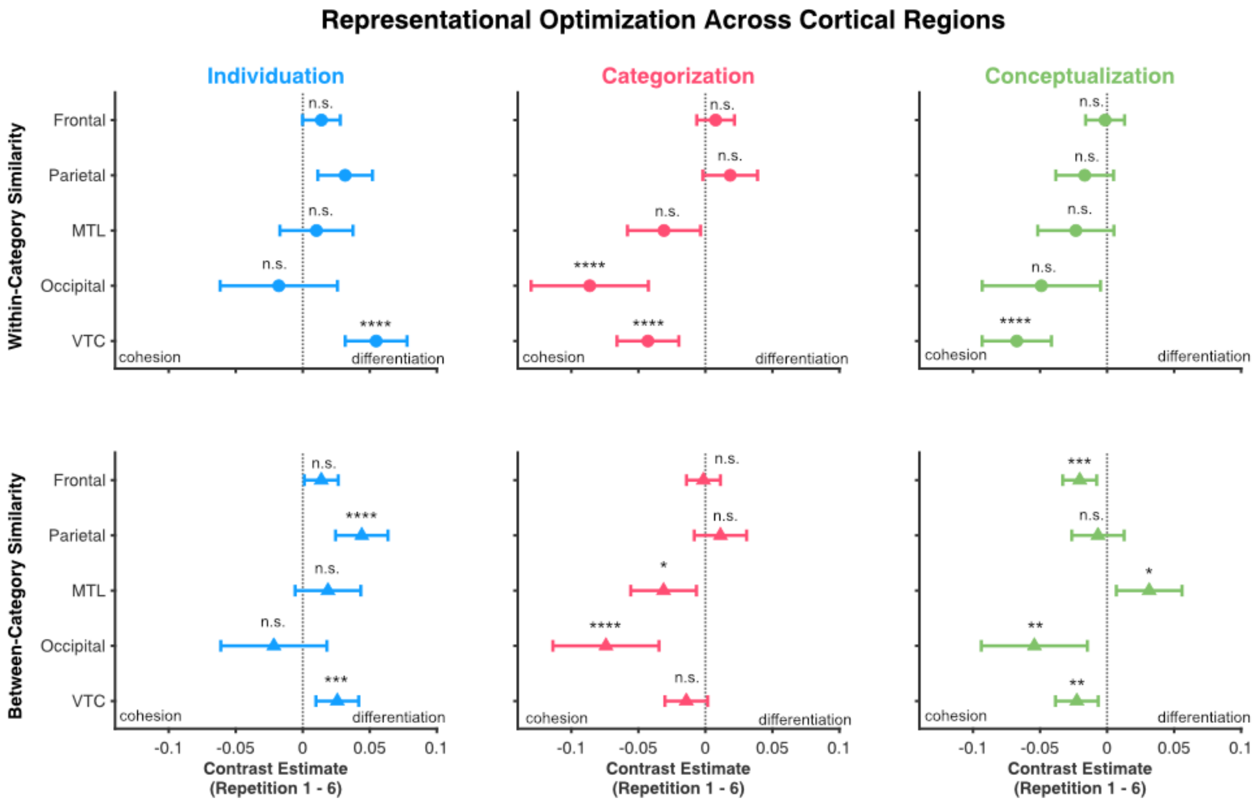


#### Fig. S4: Representational optimization across brain regions

Repetition 1–6 contrast estimates are shown for within-category (top; circles) and between-category similarity (bottom; triangles) across tasks and regions. Negative values indicate increasing similarity (cohesion), whereas positive values indicate decreasing similarity (differentiation). Error bars show 95% confidence intervals (*$p < 0.05$, **$p < 0.01$, ***$p < 0.001$, ****$p < 0.0001$; n.s., not significant). All statistics shown here are in the legend of Fig. 3 and this is a rearrangement of the data to facilitate the comparison of brain regions.

### Sections

#### 1. Validation of temporal pattern separation measure

We determined whether there was higher within and/or between category similarity between time courses after stimulus onset than during an equally long baseline period immediately preceding the stimuli. This would be expected if representations assumed a well-structured, low-dimensional geometry upon stimulus presentation[1]. Indeed, we find that all areas show an increase in within and between category similarities over the baseline (main effect of time point, all $p<0.0359$), with the exception of within category similarities in the MTL that do not reach statistical significance ($F(1, 341)=0.7932$, $p=0.3737$).

#### 2. Spatial population-level representational geometry

The temporal pattern-separation analysis captures information encoded in the temporal evolution of responses at individual electrodes. However, representational information may also be encoded in the spatial pattern of activity over electrodes within a cortical region. We therefore performed a complementary population-level RSA[2] on the spatial pattern of responses to test representations encoded in this dimension were immediately task-tailored at repetition 1 or became progressively optimized over successive repetitions.

To specifically assess the spatial population code, rather than temporal response patterns within individual electrodes, we averaged each electrode's activity across the same stimulus epoch used in the temporal RSA (0–1 s). To ensure direct comparability to the existing analysis of the temporal domain, we use the same analytic approach previously reported in the manuscript. Spatial RSA was computed separately for each participant and anatomical cluster (minimum number of electrodes=4) and multivariate noise normalization was applied separately for each similarity matrix. To this end, a covariance matrix was estimated from activity pooled across trials and time points, treating trial-by-time samples as observations and electrodes as variables. The covariance estimate was regularized using Ledoit-Wolf shrinkage[3], and its inverse square root was applied to whiten the multielectrode patterns, thereby rescaling and decorrelating electrode responses before RSA[4,5]. For each participant, region, task, and repetition, the whitened responses across electrodes formed multivariate population vectors. Cosine similarity was calculated for every stimulus pair and retained at the image-pair level. Within-category comparisons comprised face-face and house-house pairs, whereas between-category comparisons comprised face-house pairs.

To test for whether instantaneous transformation occurs through spatially distributed population-level codes, we calculated two task-tailored transformation scores at repetition 1 (similar to the main analysis). The within-category score contrasted the mean similarity during categorization and conceptualization against individuation. The between-category score contrasted conceptualization against the mean of categorization and individuation. Positive values on both dimensions indicate the geometry predicted to be optimal for the current tasks. These scores were tested against zero using one-sided t-tests. At repetition 1, within-category transformation scores were nonsignificant in frontal cortex, $t(10) = 0.27$, $p = 0.396$; parietal cortex, $t(8) = -0.15$, $p = 0.556$; MTL, $t(9) = 0.22$, $p = 0.417$; occipital cortex, $t(6) = -0.40$, $p = 0.648$; and VTC, $t(13) = 0.48$, $p = 0.321$. Between-category scores were also nonsignificant in frontal cortex, $t(9) = -1.36$, $p = 0.897$; parietal cortex, $t(7) = 0.89$, $p = 0.201$; MTL, $t(7) = -0.73$, $p = 0.757$; occipital cortex, $t(5) = 0.44$, $p = 0.339$; and VTC, $t(11) = -1.24$, $p = 0.879$.

We additionally fitted separate linear mixed-effects models to within- and between-category similarities for each region at repetition 1. Task and electrode count were included as fixed effects, with participant and image pair as random intercepts. Electrode count was log-transformed and z-scored to control for variation in population dimensionality. One-sided model-based contrasts were evaluated in the predicted directions and the p-values were combined using Stouffer's method (similar to the temporal RSA).

The mixed-effects analysis confirmed the aforementioned result: no region showed a statistically significant task-tailored transformation after combining the predicted within- and between-category contrasts using Stouffer's method (frontal: $z = -2.461$, $p = 0.9931$; parietal: $z = 0.822$, $p = 0.2055$; MTL: $z = 0.407$, $p = 0.3421$; occipital: $z = -0.494$, $p = 0.6893$; VTC: $z = -1.340$, $p = 0.9099$). Thus, although participants successfully performed all three tasks from repetition 1 onwards, we found no statistically significant evidence that spatial population representations were immediately transformed into task-tailored geometries.

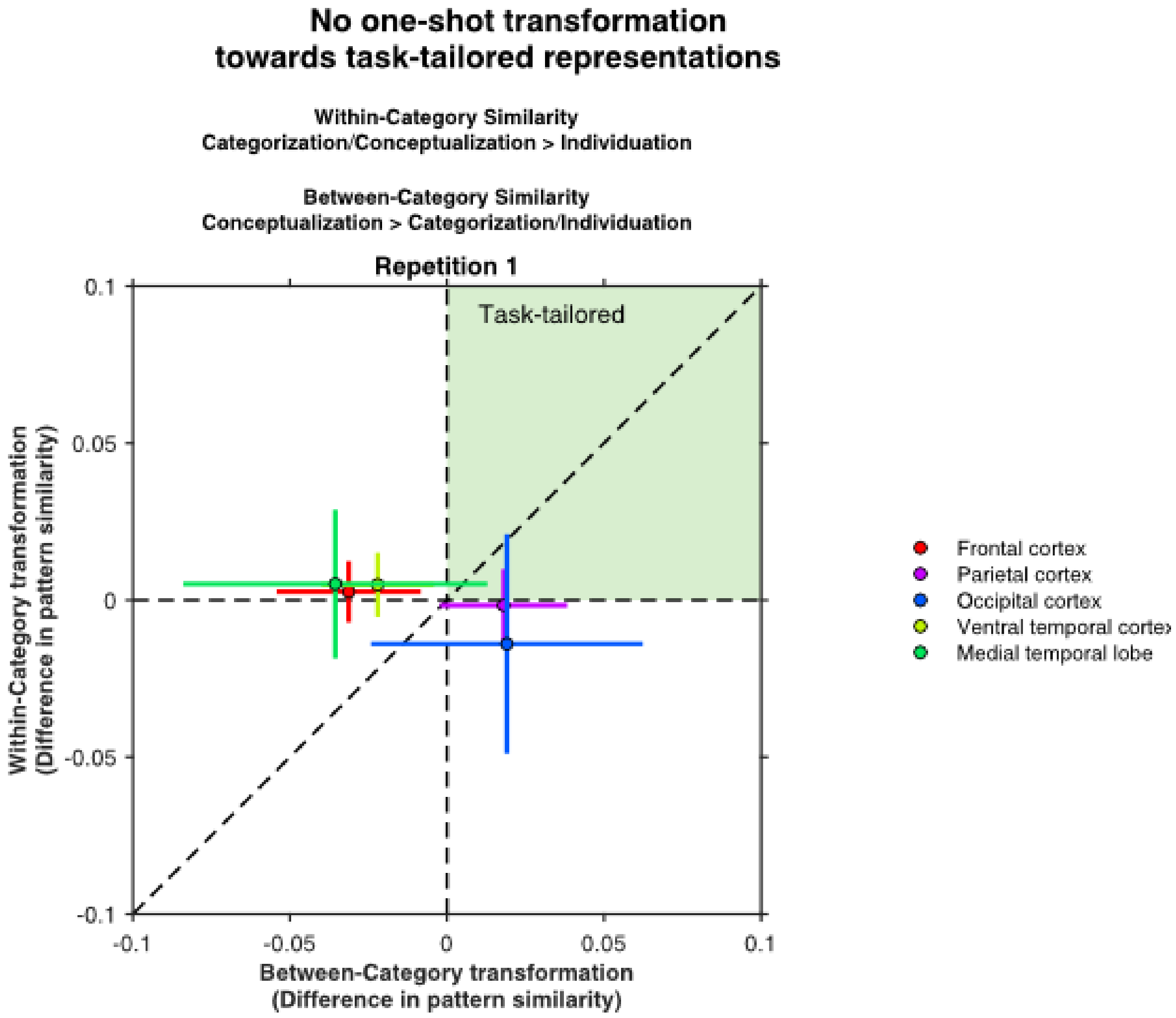


Fig. S5: No one-shot transformation towards task-tailored representations

The 2-D plot shows task-tailored similarity contrasts based on the spatial pattern of activity across electrodes in each ROI: within-category similarity (x-axis; categorization/conceptualization > individuation) and between-category similarity (y-axis; conceptualization > categorization/individuation). Positive values on both axes indicate task-tailored representations. No region showed statistically significant optimization at repetition 1 (one-sided t-tests, all $p > 0.20$).

To test gradual optimization, separate mixed-effects models were fitted to within- and between-category image-pair similarities across repetitions. Follow-up models within each region tested the task × repetition interaction while controlling for electrode count and including participant and image pair as random intercepts. Observations with residual z-scores exceeding ±3.5 were excluded at the image-pair level before refitting. Repetition 1 versus repetition 6 contrasts were Holm-corrected across the three tasks within each region and similarity type.

For within-category similarity, the task × repetition interaction was nonsignificant in frontal cortex, $F(10, 22{,}814.03) = 0.67$, $p = 0.756$; parietal cortex, $F(10, 16{,}657.19) = 1.15$, $p = 0.320$; and MTL, $F(10, 20{,}149.36) = 0.41$, $p = 0.943$. Occipital cortex showed a significant interaction, $F(10, 13{,}422.63) = 3.70$, $p < 0.001$, but none of the task-specific repetition contrasts survived Holm correction, all $p > 0.064$. VTC also showed a significant interaction, $F(10, 27{,}333.51) = 2.11$, $p = 0.020$, with significant differentiation during individuation, $t(27{,}171.69) = 4.04$, $p < 0.001$; the remaining contrasts were nonsignificant, all $p > 0.171$.

For between-category similarity, the interaction was nonsignificant in frontal cortex, $F(10, 23{,}533.19) = 1.69$, $p = 0.077$; MTL, $F(10, 20{,}983.36) = 1.00$, $p = 0.443$; and occipital cortex, $F(10, 14{,}040.51) = 1.27$, $p = 0.242$. Parietal cortex showed a significant interaction, $F(10, 17{,}143.29) = 1.92$, $p = 0.038$, with decreasing similarity, i.e., differentiation, during individuation, $t(17{,}071.66) = 3.48$, $p = 0.001$; the other contrasts were nonsignificant, all $p > 0.299$. VTC also showed a significant interaction, $F(10, 28{,}348.65) = 1.89$, $p = 0.041$, with differentiation during individuation, $t(28{,}220.68) = 4.54$, $p < 0.001$; the remaining contrasts were nonsignificant, all $p > 0.453$.

Hence, we did not find evidence that the spatial population code showed parallel optimization for all three tasks. Instead, it primarily captured differentiation during individuation, complementing the temporal code in VTC that showed optimization across all three tasks. The progressive differentiation during individuation may reflect the emergence of a contrastive code[6,7]. This is consistent with proposals that instance-level separability is a key objective of high-level visual representations[8,9]. Accordingly, differentiation in VTC and parietal cortex may represent task-driven optimization for distinguishing individual stimuli.

Overall, the population-vector analysis captures representational information distributed across electrodes. We found no statistically significant evidence that spatial population geometry rapidly adopts task-tailored configurations for all three tasks; instead, it gradually differentiated stimuli during individuation. This suggests that temporal and spatial response patterns capture complementary dimensions of gradual task-dependent representational optimization.

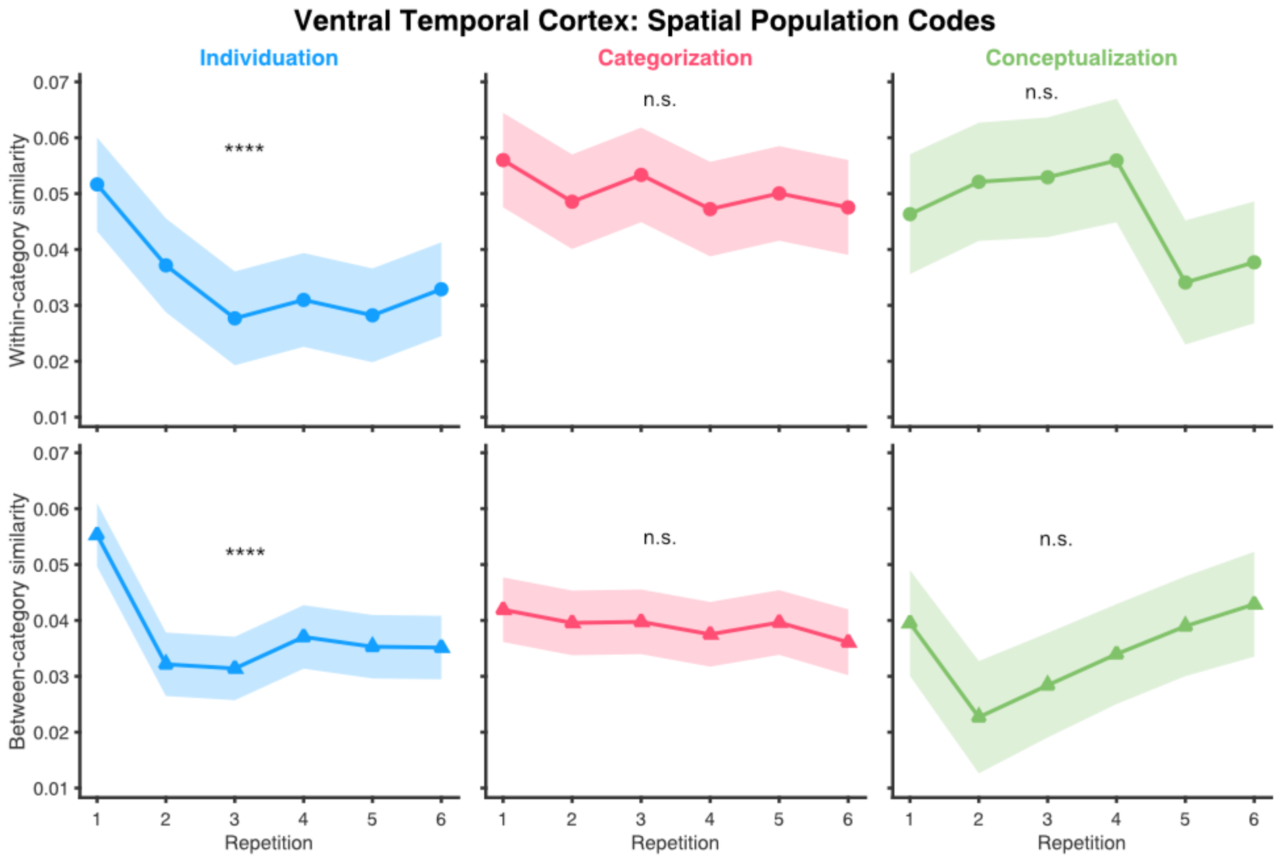


**Fig. S6 Spatial population codes in ventral temporal cortex**

Estimated marginal means for within-category (top) and between-category similarity (bottom) across repetitions and tasks. Task × repetition interactions were significant for within-category similarity, $F(10, 27333.51) = 2.11$, $p = 0.020$, and between-category similarity, $F(10, 28348.65) = 1.89$, $p = 0.041$. Repetition 1–6 contrasts were significant only for individuation (within: $t(27171.69) = 4.04$, $p < 0.0001$; between: $t(28220.68) = 4.54$, $p < 0.0001$), but not categorization (within: $p = 0.085$; between: $p = 0.227$) or conceptualization (within: $p = 0.415$; between: $p = 0.766$). This indicates that spatial population codes in ventral temporal cortex exhibit differentiation of stimuli, both within and between-category for the individuation task, but no other statistically significant optimizations. Shading denotes ±SE, **** $p < 0.0001$, n.s.: non-significant.

##### 3. VTC representational effects remain robust after matching electrode counts across regions

VTC contained more visually responsive electrodes than the other regions (see Supplementary Table 1), potentially increasing sensitivity to representational changes. To make sure that the VTC effects do not merely reflect denser sampling, we repeated the analysis after randomly subsampling VTC to 66 electrodes, corresponding to the average number of electrodes across non-VTC regions. Subsampling was repeated 100 times separately for within- and between-category similarity. For each subsample, we fitted the original VTC mixed-effects model: similarity ~ repetition × task + (1 | subject/electrode). We extracted the repetition 1 minus repetition 6 contrast for each task. Positive contrasts indicate decreasing similarity across repetitions, or differentiation, whereas negative contrasts indicate increasing similarity, or cohesion. Stability was assessed from the median bootstrap t-statistic, its 2.5th-97.5th percentile interval, and an empirical two-sided bootstrap p-value.

For within-category similarity, the original pattern was preserved: differentiation during individuation (median $t = 2.35$, 95% bootstrap interval [1.40, 3.31], $p = 0.01$) and cohesion during categorization (median $t = -1.68$, interval [−2.98, −0.19], $p = 0.04$) and conceptualization (median $t = -3.37$, interval [−4.96, −1.99], $p = 0.01$). Between-category similarity also showed differentiation during individuation (median $t = 2.19$, interval [0.93, 3.34], $p = 0.01$) and cohesion during conceptualization (median $t = -1.93$, interval [−3.41, −0.40], $p = 0.01$). Categorization showed no significant change (median $t = -1.21$, interval [−2.31, 0.01], $p = 0.06$). Thus, matching VTC electrode coverage to the average non-VTC coverage preserved the original task-dependent pattern, indicating that the distinct VTC profile was not simply attributable to denser electrode sampling.

#### 4. Neural amplitude changes for multiple tasks

Previous literature suggests repetitions of stimuli do not only entail changes in representational geometry but also changes the neural activity amplitudes. This is the well-known repetition suppression effect, which has been observed in occipital, temporal, and frontal cortex[10,11] as well as in the MTL[12] across a variety of tasks and stimuli. Such decreases in evoked activity are thought to contribute to optimizing for energy efficiency[13,14]. Some accounts imply that repetition suppression reflects adaptation or neural fatigue[15], whereas others suggest that it is a consequence of sensory predictions and hence reflects minimizations of prediction errors[13,16]. If decreases in neural activity were simply due to adaptation, then one would not expect these changes to be differential based on task demands – especially if visually dissimilar stimuli repeat, as is the case in the conceptualization task. We thus set out to determine which areas show differential repetition suppression in the tasks at hand.

Given that we observed flexible representational optimization in ventral temporal cortex, we ask whether repetition suppression also occurs differentially here. We find that the ventral temporal cortex shows an interaction between task and repetition (F(10, 4568)=2.1396, p=0.0186): amplitudes are reduced between the first and the sixth repetition in the individuation (repetition 1 vs. 6, t(4568)=2.809, p=0.01; linear trend, t(4568)=-3.151, p=0.0065; quadratic trend, t(4568)=2.692, p=0.0214) and categorization tasks (repetition 1 vs. 6, t(4568)=3.579, p=0.001; quadratic trend, t(4568)=-3.342, p=0.0034), but not in the conceptual task (repetition 1 vs. 6, t(4568)= 1.233, p=0.2177; Fig. S7) (multiple-comparisons correction done for 3 tasks and 4 trends, respectively). This suggests that task-specific repetition suppression occurs in the ventral temporal cortex but not for all three tasks. Hence, even though representational changes flexibly optimize all three tasks in ventral temporal cortex, amplitude changes here do not reflect flexible changes, particularly when abstract concepts are to be learned.

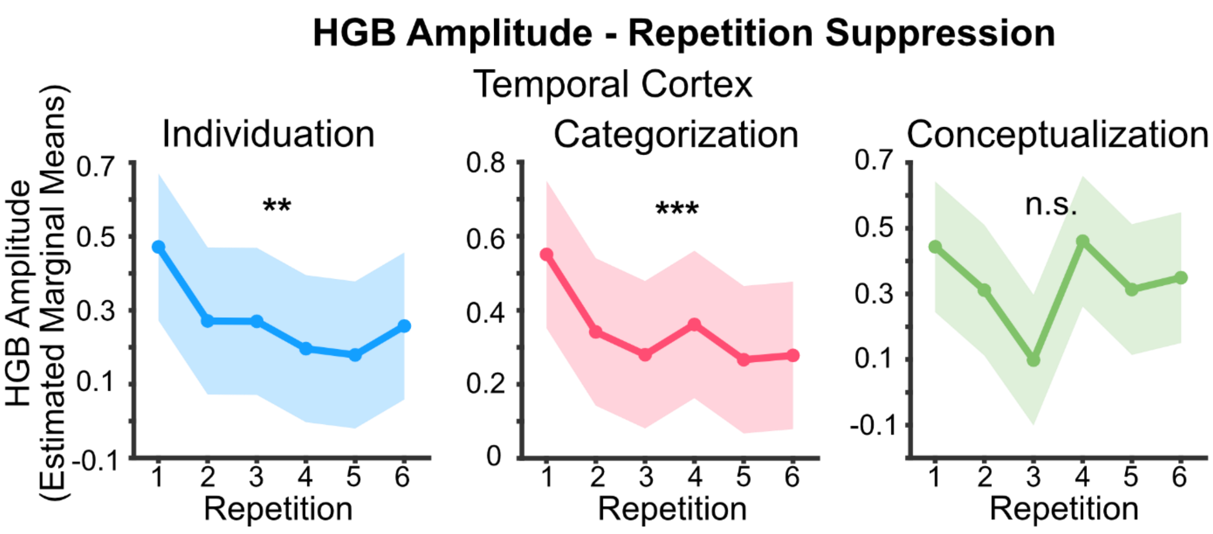


**<u>Fig S7: High-gamma band amplitude repetition suppression for multiple tasks in the ventral temporal cortex</u>**

Repetition suppression is assessed by computing decreases in amplitude of the HGB. Ventral temporal cortex exhibits repetition suppression in two out of the three tasks (interaction task × repetition, F(10,4568)=2.1396, p=0.0186): in the individuation task (repetition 1 vs. 6, t(4568)=2.809, p=0.01) and the categorization tasks (repetition 1 vs. 6, t(4568)=3.579, p=0.001), but not the conceptualization task (repetition 1 vs. 6, t(4568)=1.233, p=0.2177). All p-values are corrected for multiple comparisons for the number of tasks. * p<0.05 ; ** p<0.025; *** p<0.005; **** p<0.0005; n.s. not significant; asterisks denote statistical test done between repetition 1 and 6.

In occipital cortex, we find no statistically significant modulation of repetition suppression based on the specific task (task × repetition interaction, F(10,1488)=0.8121, p=0.6171). Yet, we find a modulation of amplitudes by the different tasks (main effect of task, F(2,1488)=8.3411, p=0.0002), as well as overall repetition suppression (main effect of repetition, F(5,1488)=3.6043, p=0.0030). Amplitudes are higher in the conceptual than in the categorical task (t(1489)=3.038, p=0.0073) but do not differ significantly between conceptualization and individuation (t(1489)=1.116, p=0.2645) nor between individuation and categorization tasks (t(1489)=1.922, p=0.1096) (multiple-comparisons correction done for 3 task comparisons). Amplitudes vary in a quadratic fashion (t(1488)=3.829, p=0.0005, multiple-comparisons correction done for 4 trends) over repetitions, but this does not depend upon the task. This indicates that the overall amplitude in occipital cortex is differentially modulated by the tasks and they are also continuously reduced, but that the latter likely constitutes a general effect that does not specifically optimize individual tasks or contribute to flexible multitask learning.

Lastly, MTL, frontal, and parietal cortex only show task modulations but no statistically significant optimization of amplitudes over repetitions (Fig. S8). Overall, this suggests that while different areas show modulation of amplitudes based on the task, flexible learning in terms of a task-specific optimization of stimulus-evoked activity levels only occurs in ventral temporal cortex, but not for all the three tasks.

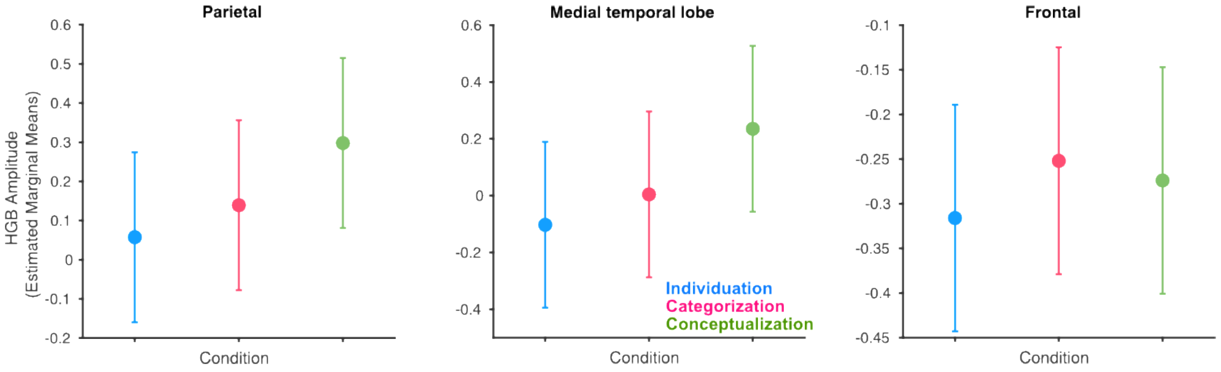


**Fig. S8: Differential task effects in terms of High-Gamma Band (HGB) amplitudes**

Depicted are normalized HGB amplitude's estimated marginal means, along with its model-based standard error. Blue indicates individuation task (In), pink indicates categorization task (Ca) and green indicates conceptualization task (Co). Parietal: we found a main effect of task (F(2,2048)=21.2983, p=7.001e-10). Task adjustment in terms of amplitude was as follows: In=Ca<Co (In=Ca: t(2048)=-2.183, p=0.0291; Co>Ca: t(2048)=4.235, p<0.0001; Co>In: t(2048)=6.418, p<0.0001). Medial temporal lobe: we found a main effect of task (F(2, 1138)=14.9698, p=3.826e-07). Task adjustment in terms of amplitude was as follows: In=Ca<Co (In=Ca: t(1138)=-1.697, p=0.0900; Co>Ca: t(1138)=3.657, p=0.0005; Co>In: t(1138)=5.353, p<0.0001). Frontal: we found a main effect of task (F(2, 4008)=3.1532, p=0.0428). Task adjustment in terms of amplitude was as follows: (In<Ca: t(4008)=-2.467, p=0.0410; Co=Ca: t(4008)=-0.826, p=0.4088; Co=In: t(4008)=1.641, p=0.2018).

So far, this suggests that amplitude changes are more indicative of adaptation than differential changes due to task demands. However, it is possible that task demands induce repetition suppression only in subjects exhibiting high task accuracy. To test this possibility, we investigate whether repetition suppression in ventral temporal cortex, specifically in the conceptualization task, is contingent upon accuracy of the subjects. Although we find an interaction effect of group and repetition (group × repetition $F(5,1431)=2.7736$, $p=0.0168$), there is no statistically significant repetition suppression in either of the groups (both $p>0.19$) and we find no significant evidence that differences between the repetitions depend on accuracy ($t(1431)=-1.652$, $p=0.0987$). Hence, there is no demonstrable evidence for task-dependent repetition suppression in ventral temporal cortex in the conceptualization task, even when considering behavioral accuracy. Overall, the above results indicate that repetition suppression in ventral temporal cortex does not occur in a differential manner for the three tasks tested here, particularly when stimuli are visually distinct from each other.

#### 5. Dependence of the result on pre-selection of visually responsive electrodes

So far, we investigated task-dependent optimization using analyses of representational geometry of visually responsive electrodes as identified using an independent functional localizer. However, representational geometry can be shaped not only by strongly stimulus-responsive electrodes, but also by electrodes that do not exhibit reliable selectivity or stimulus-evoked responses. Especially electrodes that do not reach a statistical threshold for visual responsiveness in the functional localizer may nevertheless contribute to population representations. We therefore asked whether the task-dependent transformations reported in the main analysis depend on restricting the electrode pool to the visually responsive electrodes identified by the independent localizer.

To determine whether our conclusions depend on visual responsiveness, we directly compared task-dependent transformations in visually responsive and non-visually responsive electrodes. Following the statistical logic of the main analyses, we fitted similar linear mixed-effects models as before, while adding visual responsivity as a categorical fixed effect.

Our first analysis examined whether task-dependent representational geometry was already present upon the first encounter with a stimulus, indicating a one-shot transformation. Because such an effect could differ between visually responsive and non-visually responsive electrodes, we first tested the task × visual-responsiveness interaction within each anatomical cluster, separately for within- and between-category similarity. As task-optimality is jointly defined by within- and between-category structure, we combined the interaction evidence across the two similarity types using Stouffer's method to obtain a single decision for each cluster. The combined interaction was not significant in frontal cortex ($z = -0.20$, $p = 0.580$), MTL ($z = 0.97$, $p = 0.166$), occipital cortex ($z = -0.07$, $p = 0.529$), or parietal cortex ($z = 1.12$, $p = 0.131$), providing no evidence that task-dependent representations differed according to visual responsiveness in these regions. Visually responsive and non-visually responsive electrodes were therefore combined for the subsequent analyses in these clusters. In contrast, VTC showed a significant interaction ($z = 4.40$, $p < 0.0001$), and its visually responsive and non-visually responsive electrodes were consequently examined separately. Following the logic of the original analysis, one-shot transformation toward task-optimal geometry would require greater within-category similarity during categorization and conceptualization than during individuation, together with greater between-category similarity during conceptualization than during categorization and individuation. We combined these directed, one-sided contrasts first within each similarity type and then across similarity types using Stouffer's method. Neither the analyses combining all electrodes nor the separate analyses of visually responsive and non-visually responsive VTC electrodes revealed the predicted task-optimal geometry (all Holm-corrected $p \geq 0.288$). Thus, consistent with the findings based on visually responsive electrodes, including all electrodes provided no statistically significant evidence for an immediate task-optimal reconfiguration in any anatomical cluster.

Further, to test whether task-optimization of representational geometry occurred over repetitions in each ROI, linear mixed effects models were fitted using repetition, task, visual responsivity, and their interactions as fixed effects, with electrodes nested within participants as random intercepts in a similar manner to the main manuscript. A non-significant interaction provides no evidence that task-dependent representational optimization differ according to visual responsiveness in that region of interest; in these cases, visually responsive and non-visually responsive electrodes were combined to estimate their common representational optimization. A significant interaction indicates that visual responsiveness modulates representational change over repetitions. In these cases, pooling the two populations could obscure distinct transformations, and we therefore additionally characterized the non-visually responsive electrodes separately.

For between-category similarity, the interaction was significant in VTC, $F(10, 7055) = 5.8885$, $p < 0.0001$, and frontal cortex, $F(10, 5610) = 2.5149$, $p = 0.0051$, but not in parietal cortex, $F(10, 3128) = 0.8088$, $p = 0.6202$, MTL, $F(10, 1445) = 0.9794$, $p = 0.4593$, or occipital cortex, $F(10, 833) = 0.6129$, $p = 0.8037$. For within-category similarity, this interaction was significant in VTC, $F(10, 13131) = 4.2208$, $p < 0.0001$, and parietal cortex, $F(10, 5555) = 1.9225$, $p = 0.0377$. It was not significant in frontal cortex, $F(10, 10025) = 1.3779$, $p = 0.1835$, MTL, $F(10, 2715) = 1.7372$, $p = 0.0671$, or occipital cortex, $F(10, 1807.01)=1.1811$, $p=.2988$.

In occipital cortex, neither visual-responsiveness interaction was statistically significant. The combined analysis showed no task x repetition interaction for between-category similarity, $F(10, 833) = 1.4387$, $p = 0.1583$, or within-category similarity, $F(10, 1824) = 0.9083$, $p = 0.5245$. The repetition 1 versus 6 contrasts likewise showed no significant task-specific changes, all $p > 0.14$. Thus, consistent with the analysis of only visually responsive electrodes, occipital cortex showed no evidence of gradual task-tailored representational optimization.

In MTL, we found no statistically significant evidence that visual responsiveness modulated either between- or within-category transformations, allowing inclusive analyses. Between-category similarity showed a significant task x repetition interaction, $F(10, 1411) = 4.5163$, $p < 0.0001$. Similarity decreased during ad hoc concept formation, indicating differentiation, $t(1411) = 4.0881$, $p = 0.0001$, and during individuation, $t(1411) = 2.3654$, $p = 0.0363$. In contrast, similarity increased during categorization, indicating cohesion, $t(1411) = -2.2031$, $p = 0.0363$. Within-category similarity also showed a significant task x repetition interaction, $F(10, 2732) = 3.5714$, $p = 0.0001$. However, none of the corrected repetition 1 versus 6 contrasts was significant: ad hoc concept formation, $t(2732) = 1.0755$, $p = 0.2822$; individuation, $t(2732) = 2.0128$, $p = 0.1327$; or categorization, $t(2732) = -1.7440$, $p = 0.1625$. MTL representations therefore changed over repetitions, but these transformations did not consistently follow the task-optimal geometry. This is consistent with the analysis of visually responsive electrodes, in which MTL transformations ran counter to the representational structure required by the task.

In frontal cortex, visual responsiveness modulated the between-category transformation. However, non-visually responsive electrodes showed no significant task x repetition interaction, $F(10, 3621) = 1.4661$, $p = 0.1455$, and therefore no reliable task-dependent cohesion or differentiation (individuation, $t(3621) = 1.9953$, $p = 0.0922$, or categorization, $t(3621) = -0.4960$, $p = 0.6200$, ad hoc conceptualization $t(3621) = 2.3811$, $p = 0.0519$). Thus, the between-category cohesion supporting ad hoc concept formation in the primary analysis was specific to visually responsive frontal electrodes. For within-category similarity, visual responsiveness did not significantly modulate the transformation, permitting an analysis combining non-visually and visually responsive electrodes. This analysis showed a significant task x repetition interaction, $F(10, 10077) = 2.9716$, $p = 0.0010$, including differentiation during individuation, $t(10077) = 3.0572$, $p = 0.0045$, and cohesion during ad hoc concept formation, t(10078)

$= -3.5222$, $p = 0.0013$. Categorization did not produce a significant change, $t(10077) = 1.2833$, $p = 0.1994$. Frontal cortex therefore contributed to representational cohesion during ad hoc concept formation and within-category differentiation during individuation, but did not exhibit task-tailored transformations across all three tasks, consistent with the analysis of visually responsive electrodes.

In parietal cortex, visual responsiveness did not significantly modulate the between-category optimization. The combined analysis showed no significant task x repetition interaction, $F(10, 3162) = 1.2861$, $p = 0.2321$, and therefore no reliable task-dependent cohesion or differentiation. Consistent with this result, the repetition 1 versus 6 contrasts were not significant for ad hoc concept formation, $t(3162) = -1.2782$, $p = 0.4026$, individuation, $t(3162) = 2.3151$, $p = 0.0620$, or categorization, $t(3162) = 1.0184$, $p = 0.4026$. Non-visually responsive electrodes showed no significant task x repetition interaction, $F(10, 3692) = 1.3052$, $p = 0.2213$, and no task showed significant task-dependent optimization between repetitions 1 and 6: individuation, $t(3692) = 2.3512$, $p = 0.0563$; or categorization, $t(3692) = 1.1521$, $p = 0.4128$ or ad hoc concept formation, $t(3694) = -1.2638$, $p = 0.4128$. Thus, the task-tailored individuation effect reported in parietal cortex was expressed most clearly among visually responsive electrodes and was not reliably present in the non-visually responsive population.

Finally, visual responsiveness modulated both between- and within-category transformations in VTC. Among non-visually responsive electrodes, between-category similarity showed a significant task x repetition interaction, $F(10, 4896) = 7.5448$, $p < 0.0001$. Ad hoc concept formation produced significant differentiation, $t(4896) = 4.0675$, $p = 0.0001$, which was opposite to the task-tailored cohesion observed among visually responsive VTC electrodes. Neither individuation, $t(4896) = 1.7192$, $p = 0.1713$, nor categorization, $t(4896) = -1.6394$, $p = 0.1713$, showed a significant repetition 1 versus 6 transformation. Within-category similarity also showed a significant task x repetition interaction, $F(10, 8418) = 2.8433$, $p = 0.0016$. However, none of the corrected repetition 1 versus 6 contrasts was significant: ad hoc concept formation, $t(8418) = -2.0853$, $p = 0.1112$; individuation, $t(8418) = 1.7094$, $p = 0.1748$; or categorization, $t(8418) = -1.4074$, $p = 0.1748$. Thus, although non-visually responsive VTC electrodes changed over repetitions, they did not show the complete task-tailored profile expressed by visually responsive VTC electrodes. Contrasting changes in the representational structure of stimulus-responsive and non-responsive neuronal populations have previously been shown to contribute to the stability of sensory representations[26]. The functional significance of the opposing VTC effect observed here remains to be determined.

Taken together, these analyses clarify how visual responsiveness affects the expression of task-dependent representational change. VTC remains the only region showing robust task-tailored optimization across all three tasks, and this complete profile is expressed by independently identified visually responsive electrodes. MTL continues to show repetition-dependent transformations that do not consistently align with task-optimal geometry, occipital cortex remains largely non-task-tailored, and frontal and parietal effects depend more strongly on visual responsiveness. The combined analyses did not reveal an alternative task-tailored organization among non-visually responsive electrodes that could account for the primary findings.

Visual responsiveness therefore does not appear to artifactually create the reported representational geometry. Rather, independent functional localization identifies the electrodes in which task-relevant transformations are expressed most reliably, consistent with attention and predictive-coding accounts in which the most informative neural populations contribute most strongly to task-dependent changes[17,18]. At the same time, the reduced and occasionally opposing effects among non-visually responsive electrodes illustrate how adding measurements with weaker stimulus-related signals can reduce sensitivity of the analyses[19,20]. This makes the interpretation of results challenging, and motivates feature-selection, ridge-based penalties etc. on non-responsive and noisy neurons/voxels/electrodes in the single neuron/fMRI/iEEG literature[17,21,22,23,24,25].

#### 6. Split-Half Reliability of Localizer-Based Temporal Pattern Similarity

Representational similarity measures are based on correlations between neural activity patterns which can be affected by differences in trial-to-trial reliability, signal stability, or measurement noise in a certain brain area. This noise structure could hypothetically account for the regional differences, potentially reflecting differences in the reliability of the similarity estimate rather than a genuine difference in neural representational structure. To address this concern, we performed a split-half stability analysis using the independent localizer data.

For each electrode, we extracted localizer HGB responses to faces and houses and computed temporal pattern similarity using the same logic as in the main analysis. Within-category similarity was computed separately for faces and houses, and between-category similarity was computed for face-house comparisons. Correlation values were averaged after Fisher's z-transformation. To estimate the reliability of these similarity values, we randomly split localizer trials into two independent halves 50 times for each electrode and similarity type, requiring at least four trials per half. For each split, temporal similarity was recomputed separately in each half. We then quantified instability as the absolute difference between the two split-half estimates, |split 1 - split 2|, and averaged this value across the 50 iterations. Lower values therefore indicate more stable, less noise-sensitive similarity estimates.

We modeled split-half instability separately for within-category and between-category similarity using linear mixed-effects models with cluster and visual responsiveness as predictors:

abs_splitdiff_mean ~ cluster * visually_responsive + (1 | subject/electrode)

Similar to the main analysis, when the nested random-effects structure did not converge, alternative random-effects structures were compared, and the best-fitting convergent model was retained, following the same model-selection logic used in the main analyses. Omnibus effects were evaluated using Type III ANOVA with Kenward-Roger degrees of freedom, and post hoc cluster comparisons were Holm-corrected.

For within-category similarity, there was a significant main effect of cluster, $F(4, 1108.27) = 15.67$, $p = 1.76 \times 10^{-12}$, a significant main effect of visual responsiveness, $F(1, 1103.95) = 164.86$, $p = 2.85 \times 10^{-35}$, and a significant cluster-by-visual-responsiveness interaction, $F(4, 1103.35) = 5.14$, $p = 4.19 \times 10^{-4}$. Holm-corrected post hoc comparisons showed that this cluster effect was driven by greater instability in occipital electrodes relative to all other clusters: occipital versus frontal, $p = 3.73 \times 10^{-13}$; occipital versus parietal, $p = 2.08 \times 10^{-12}$; occipital versus VTC, $p = 8.01 \times 10^{-12}$; and occipital versus MTL, $p = 3.22 \times 10^{-11}$. All remaining non-occipital pairwise comparisons were not significant after Holm correction, with all $p>0.99$.

For between-category similarity, there was again a significant main effect of cluster, $F(4, 1110.05) = 10.89$, $p = 1.15 \times 10^{-8}$, a significant main effect of visual responsiveness, $F(1, 1104.88) = 156.94$, $p = 9.20 \times 10^{-34}$, and a significant cluster-by-visual-responsiveness interaction, $F(4, 1103.99) = 4.19$, $p = 0.00227$. Holm-corrected post hoc comparisons showed the same qualitative pattern: occipital electrodes showed greater instability than frontal electrodes, $p = 4.77 \times 10^{-9}$; parietal electrodes, $p = 1.56 \times 10^{-8}$; VTC electrodes, $p = 9.20 \times 10^{-7}$; and MTL electrodes, $p = 1.29 \times 10^{-6}$. The remaining non-occipital comparisons were not significant after Holm correction (all $p > 0.091$).

Thus, the split-half stability analysis did not reveal broad regional differences in noise structure that could explain the main pattern-similarity findings. Instead, the only consistent regional difference was that occipital electrodes showed lower split-half stability. This effect was present for both within-category and between-category similarity and was observed across visually responsive and non-visually responsive electrodes. One possible interpretation is that early visual electrodes may be more sensitive to low-level image-dependent variability or retinotopic heterogeneity across sampled sites, although this mechanism remains speculative. Critically, VTC, MTL, parietal, and frontal electrodes showed comparable stability, supporting the interpretation that the repetition-dependent effects observed in VTC reflect representational reconfiguration rather than regional differences in the reliability of the correlation-based similarity measure.

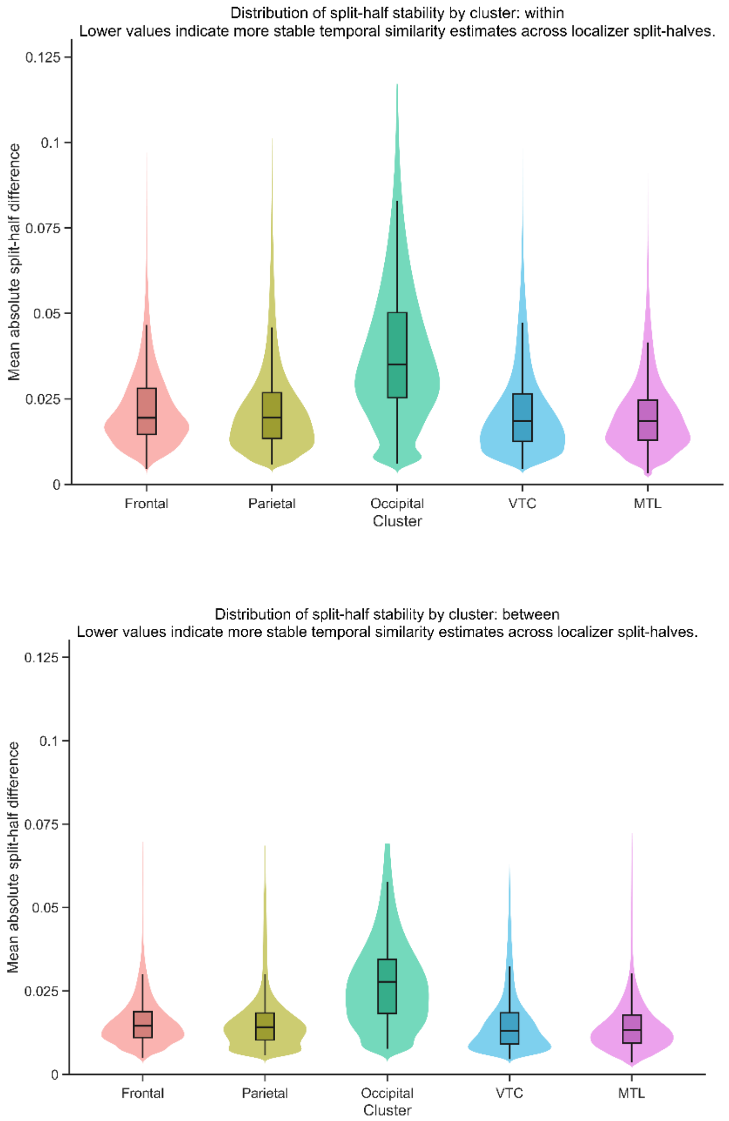


**Fig. S9: Distribution of localizer split-half stability across anatomical clusters**

Distribution of localizer split-half stability across anatomical clusters for within-category similarity (top) and between-category similarity (bottom). Violin plots show the distribution of electrode-wise mean absolute split-half differences, with embedded boxplots indicating the median and interquartile range. Lower values indicate more stable temporal similarity estimates across localizer split-halves. In both within-category and between-category analyses, occipital electrodes showed the highest split-half differences, whereas frontal, parietal, ventral temporal cortex (VTC), and medial temporal lobe (MTL) showed comparable stability.
